# Spatial transcriptomic profiling identifies lacrimal gland epithelial cell-driven mechanisms underlying autoimmunity in Sjögren’s disease

**DOI:** 10.1101/2025.11.03.686353

**Authors:** Shivali Gupta, Athanasios Ploumakis, Nikolaos Kalavros, Sharmila Masli

## Abstract

Sjögren’s disease (SjD) is a second most prevalent rheumatic disease involving autoimmune pathology of tear-producing lacrimal glands that leads to a common clinical manifestation of chronic ocular surface disease. Despite advances in understanding of SjD, lacrimal gland pathology remains incompletely understood limiting diagnosis and treatment. Here we analyze spatial transcriptomic profile of lacrimal glands from wild-type (C57Bl/6) mice and Thrombospondin (TSP)-1^-/-^ mice, a spontaneous mouse model of SjD. We uncover molecular mechanisms underlying functional loss of major epithelial cell subtypes – acinar, duct and myoepithelial cells. We identify potential early mechanisms and markers of glandular damage. By integrating spatial and cellular profiles, we uncover the presence of antigen presenting cells in the proximity of duct epithelial cells that were not described previously in lacrimal glands. We further identify role of epithelial cells as active participants in promoting or sustaining inflammation. Our findings help reveal potential molecular and cellular cues that drive periductal infiltrates containing B cells and Tfh cells that form germinal centers to facilitate local autoantibody production. Overall, our study can provide a framework for therapeutic targeting of epithelial cell types and multicellular interactions underlying autoimmune pathology.

**Significance Statement:** Tears produced by lacrimal glands are critical for protecting the eye surface by preventing tissue dryness and maintaining normal vision. Disruption of this function due to autoimmune inflammation in Sjögren’s Disease compromises the protection of the eye surface causing dryness, a condition with a potential for sight-threatening complications like infections and ulcers. Understanding cellular and molecular interactions that lead to functional loss and autoimmune inflammation of the lacrimal gland is critical for developing effective therapies. We have analyzed transcriptional profile of glandular cells in a tissue section where their morphology and interactions with surrounding cells is preserved. By comparing glands from normal and mice with Sjögren’s disease we identify molecular mechanisms that can form the basis for new therapies.

## Introduction

Sjögren’s disease (SjD) is the second most prevalent autoimmune rheumatic disease characterized primarily by chronic inflammation and dysfunction of exocrine glands like lacrimal and salivary glands. Additionally, extra-glandular manifestations involving joints, skin, lungs and nervous system due to systemic autoimmunity are observed in up to 75% patients (1). Ocular dryness or dry eye disease (DED) is one of the most common clinical manifestations of SjD resulting from lacrimal gland (LG) dysfunction and damage (2, 3). Due to insidious onset, morbidity in SjD patients increases progressively with age leading to debilitating symptoms and increased risk of mortality with extra-glandular manifestations including lymphoma (2, 4). Current management of SjD involves treatments for symptomatic relief, while there is no approved disease-modifying anti-rheumatic drug (DMARD) available (5). Despite the progress made in elaborating immunopathogenic mechanisms underlying SjD, the molecular basis of multicellular interactions in the lacrimal gland spatial architecture and tissue microenvironment that contribute to the autoimmune disease, and its progression are still not fully understood limiting therapeutic innovation in SjD.

Structurally lacrimal gland (LG) is composed of multiple lobules that contain secretory units, acini, and interlobular and intralobular duct system. Acinar epithelial cells are primary secretory cells that produce protein-rich lacrimal fluid, while the duct epithelial cells modify this primary fluid via absorption or secretion of ions thereby finetuning tear composition before delivery to the ocular surface. Contractile myoepithelial cells (MECs) wrap around both acini and ducts to facilitate expulsion of tear fluid from the gland (6). In addition to these epithelial cells, LGs also harbor innate and adaptive immune cells involved in immune defense and surveillance within the gland. Immune cells like plasma cells produce secretory IgA (sIgA) (6). Combined action of sIgA, antimicrobial factors and other proteins secreted by acinar epithelial cells creates tear fluid that is both protective and essential for maintaining the health and integrity of the ocular surface. Lacrimal gland dysfunction in SjD is known to compromise aqueous tear production and quality. While glandular damage is largely attributed to local inflammatory immune responses it is not known if and how lacrimal gland epithelial cells contribute to modulation of local immune responses and how their functional alterations collectively contribute to the pathogenesis of SjD.

In our study we used TSP-1 deficient mouse model of SjD that spontaneously develops disease with characteristic LG inflammatory infiltrates, functional loss along with ocular surface dryness and SjD-related autoantibodies as observed in human patients (7, 8). In a study designed to dissect non-immune events in NOD mouse model of SjD, microarray analysis of LG tissue from NOD/scid mice was performed (9). It is noteworthy that a search of their GEO Profiles database (10), revealed reduced expression of TSP-1 in LGs of NOD/scid mice (GEO accession GDS2177, (9)) suggesting the relevance of reduced intrinsic TSP-1 expression with disease development. Similarly, in a recent transcriptomic analysis of minor salivary glands of SjD patients, significantly reduced expression of TSP-1 was detectable (GEO accession GSE157159, (11)). This result is consistent with reduced TSP-1 expression detectable in salivary glands of patients with advanced SjD as compared to healthy controls (GEO accession GDS3940, (12)). Collectively these studies further validate clinical relevance of TSP-1 deficiency in SjD pathogenesis.

Recently several studies applied single-cell RNA-seq (scRNA-seq) to help identify diverse epithelial and immune cell populations in LGs of healthy and SjD mouse models (13-15). These studies highlight complex heterogeneity among glandular cell populations and immune cells, suggesting the need for further analysis to clarify cell-type specific contributions and molecular changes. In this study, we integrated cell-type signatures and expression programs with whole transcriptome digital spatial profiles of matched specimens of LG tissues from healthy and TSP-1 deficient SjD mouse model to elucidate cell-type specific contributions to SjD pathogenesis. We identified several distinct molecular alterations in epithelial cells that contribute to the pathogenesis. Furthermore, we discovered spatially defined interactions between epithelial cells and immune cells that support development of local adaptive immune response. These cellular mechanisms provide potential targets for therapeutic innovation in SjD.

## Results

### Mapping cell types and transcriptional programs to lacrimal gland architecture

To determine transcriptional programs of structural components of LG in the context of their spatial organization, we performed digital spatial profiling with the NanoString GeoMx mouse whole transcriptome atlas (WTA; 21,000+ genes). We used formalin-fixed paraffin-embedded (FFPE) sections of LG tissues from TSP-1^-/-^ and WT mice. Characteristic periductal and perivascular mononuclear infiltrates were detectable in H&E-stained sections of TSP-1^-/-^ but not normal WT LGs (Figure 1A). In situ hybridization was performed on these tissues, using UV-photocleavable barcode conjugated RNA probes. From selected regions of interest (ROI) mRNA counts were captured and profiled. These ROIs included acini with surrounding myoepithelial cells (MECs) or ducts and surrounding mononuclear infiltrates (Figure 1A). Using four-color immunofluorescence and morphological appearances of cell-types, custom areas of illumination (AOIs) were identified for each cell-type segment within each ROI (Figure 1B). Barcodes cleaved and collected from each AOI were quantified by sequencing. To address the previously reported heterogeneity of LG tissue we used higher sequencing depth and multiple replicates of tissue samples to reliably identify differentially expressed genes (DEGs) and help resolve signals from relatively less abundant cell types (16, 17). We analyzed total 28 LG tissue samples (supplemental figure 1) that included 4 sections from different regions of the tissue for each sample from TSP-1^-/-^ (n=4) and WT (n=3) mice. Over 27,500 transcripts obtained from 32,465 cells were sequenced and NGS data was deconvolved using scRNA-seq cell-type signatures from publicly available data set (GSE132420) as a reference. Twelve principal cell clusters were detected (Figure 1C) and their identity was validated using published genes associated with each cell type (Figure 1D). The AOIs for acinar, duct, myoepithelial and immune cells clustered appropriately by the cell-type in all selected ROIs (supplemental figure 2). Among epithelial cells, we detected most DEGs in acinar epithelial cells, followed by duct and myoepithelial cells (Table 1). In acinar epithelial cells most DEGs were upregulated, while in duct epithelial cells they were predominantly downregulated and in MECs equal proportion of DEGs were up and downregulated in TSP-1^-/-^ LGs. Further, comparisons of the major immune cell populations (T cells comprising T and NK cells, B cells, plasma cells and monocytes comprising macrophages and dendritic cells) in WT and TSP-1^-/-^ LGs revealed, increased ROIs containing B cells, plasma cells as well as T cells in TSP-1^-/-^ glands (Figure 1E).

**Table 1.** Total DEGs detected in each epithelial subtype in lacrimal gland tissues analyzed.

| Epithelial<br>cell type | Numbers of Differentially Expressed Genes (Adj p < 0.005) |  |  |  |
| --- | --- | --- | --- | --- |
|  | Total | Significantly altered | Upregulated | Downregulated |
| Acinar | 19300 | 2052 (11%) | 1182 (58%) | 870 (42%) |
| Duct | 3259 | 445 (14%) | 140 (32%) | 305 (68%) |
| Myoepithelial | 4965 | 296 (6%) | 148 (50%) | 148 (50%) |

**Figure 1:**
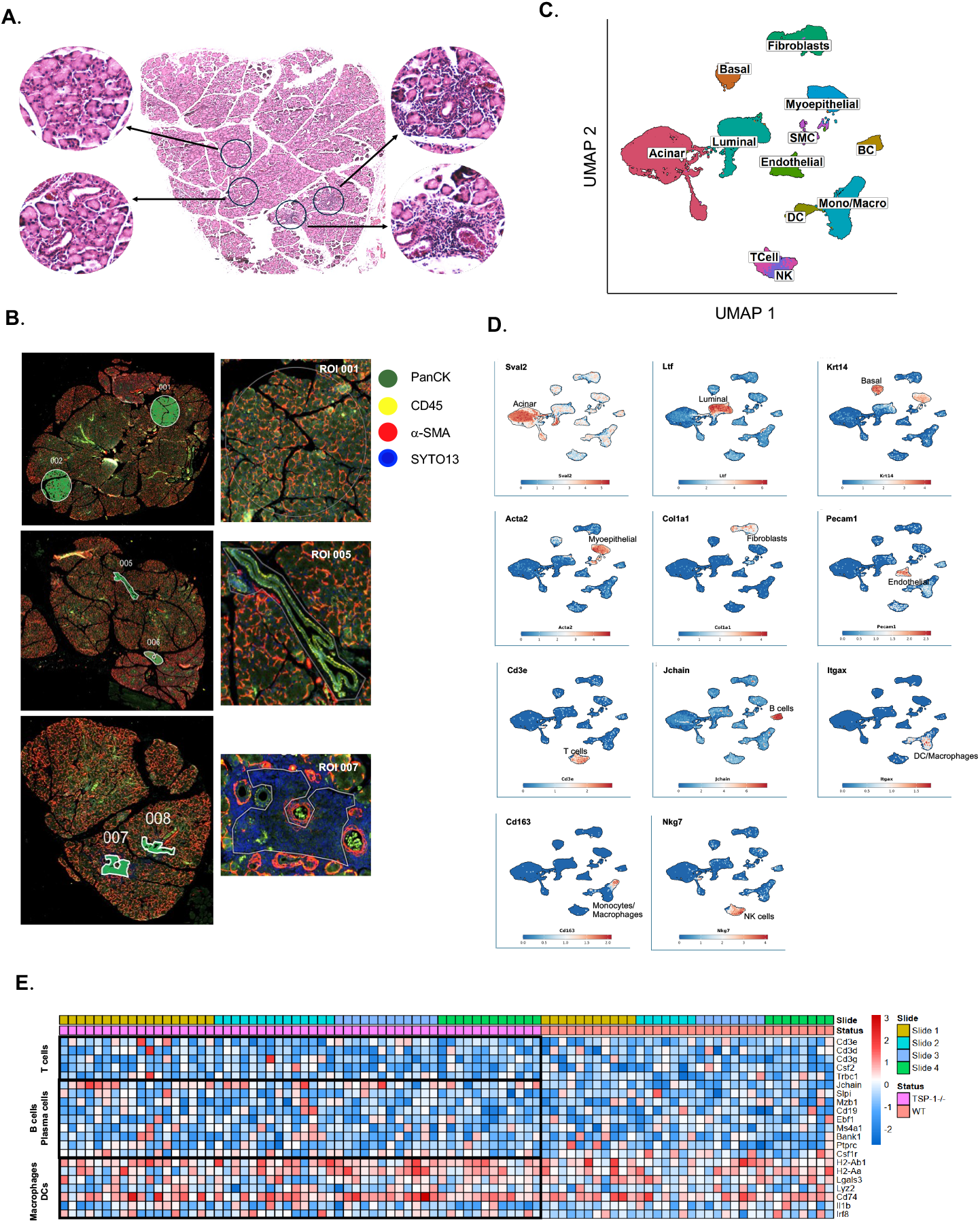
Digital spatial profiling (DSP) and Whole Transcriptome analysis (WTA) of mouse lacrimal gland tissues. **(A)** Representative hematoxylin and eosin (H&E)-stained FFPE section (5 µm thick) of LG from TSP-1^-/-^ mice showing representative areas selected for identifying region of interest (ROI) that contained acini with surrounding myoepithelial cells (upper left), ducts (lower left) and periductal and perivascular inflammatory infiltrates (upper and lower right). (**B)** Representative immunofluorescence images (GeoMx DSP) of consecutive section from the same FFPE block stained for PanCK (green), CD45 (yellow), a-SMA (red) and nuclear stain SYTO13 (blue) and ROIs. Segmentation was used to identify areas of interest (AOIs) to enrich for acinar, myoepithelial, duct and immune cells based on staining for indicated markers and morphology. (**C**) UMAP plot showing cell clusters identified in ROIs from WT and TSP-1^-/-^ LGs. **(D**) UMAP plots for indicated transcripts that identified specifically one or two clusters. (**E**) Heatmap showing major immune cell populations (T cells, B cells, plasma cells and monocytes) detected in ROIs from WT vs. TSP-1^-/-^ LGs. Color scale indicates relative expression levels.

**Figure 2.**
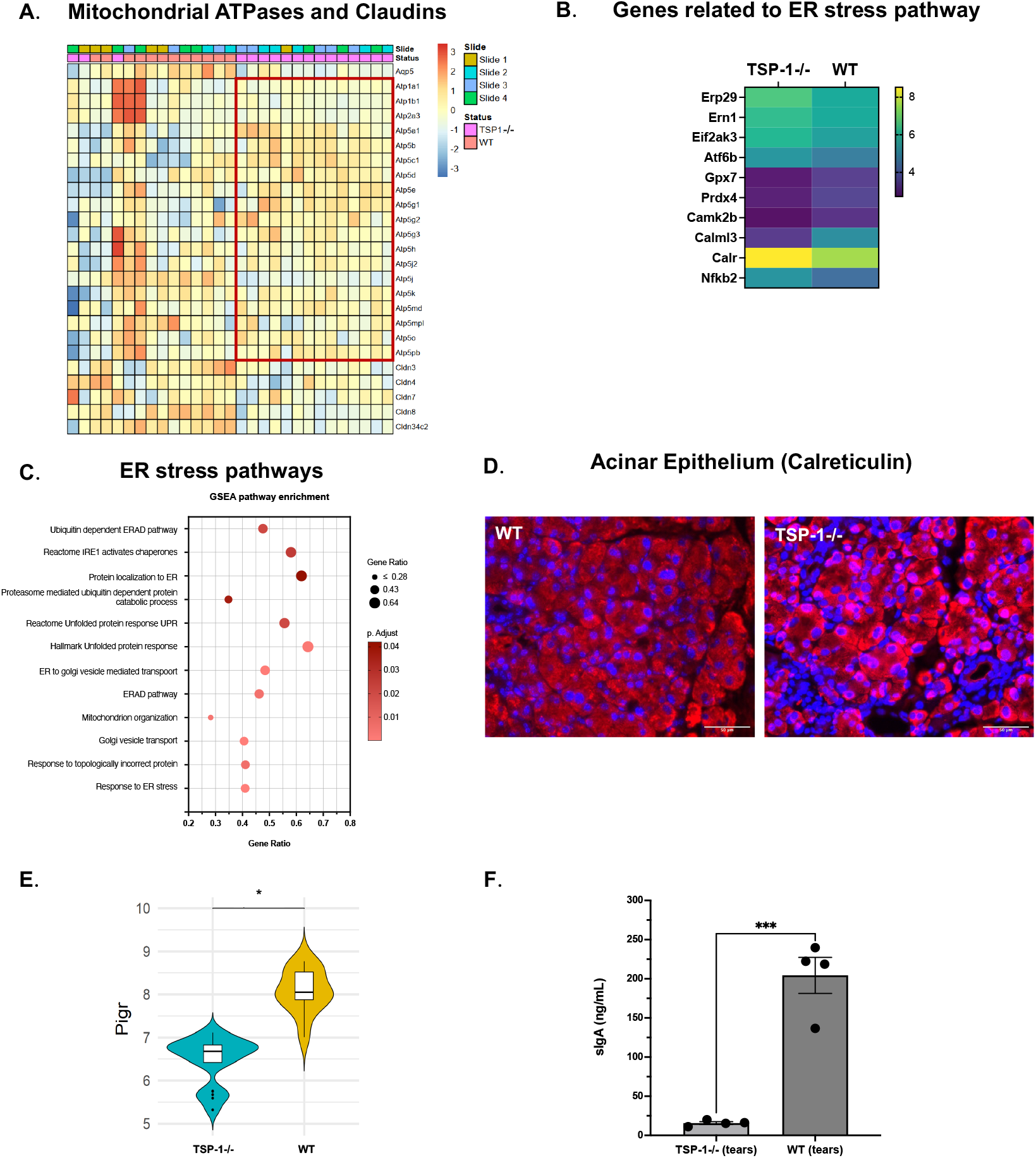
Endoplasmic reticulum (ER) stress pathways in TSP-1 deficient acinar epithelial cells and associated secretory dysfunction. **(A)** Heatmap of differential gene expression (DEG) in WT and TSP-1^-/-^ acinar epithelial cells. The red box highlights mitochondrial ATPases. **(B)** Heatmap illustrating expression changes of select genes involved in ER stress response pathways. **(C)** Pathways enriched in significantly upregulated DEGs. **(D)** Representative immunofluorescence images of Calreticulin staining (Red) in LG acinar epithelial cells. Nuclei are counterstained with DAPI (blue). **(E)** Violin plot of *Pigr* expression in WT and TSP-1^-/-^ acinar epithelial cells. **(F)** Levels of secretory IgA (sIgA) in tear samples of TSP-1^-/-^ and WT mice. (*q-value <0.05, ***p-value <0.001)

### ER stress pathways in TSP-1 deficient acinar epithelial cells disrupt their secretory function and promote autoimmune pathology

Analysis of transcriptomic profiles of acinar epithelial cells in WT and TSP-1^-/-^ LGs was performed to determine if significant differences represented potential mechanisms underlying their functional loss and autoimmunity noted in TSP-1^-/-^ mice. Increased expression of mitochondrial ATPases was detected in TSP-1^-/-^ acinar epithelial cells relative to WT control cells (Figure 2A). This increased expression of mitochondrial ATPases was accompanied by significantly downregulated expression of antioxidant enzyme genes like *Gpx7* and *Prdx4* and upregulated expression of genes associated with ER stress – *Erp29, Ern1, Eif2ak3* and *Atf6b* (Figure 2B). The presence of ER stress in TSP-1^-/-^ cells was also supported by the enrichment of pathways associated with ER stress among DEGs (Figure 2C). Furthermore, in TSP-1^-/-^ acinar cells we detected significant downregulation of *Calm3* gene (Figure 2B) that encodes epithelial-specific calcium-sensing protein like calmodulin-like protein 3 (Calml3), and *Camk2b* gene that encodes beta chain of calcium/calmodulin-dependent protein kinase II (CaMKII). Both Calml3 and CaMKII are known to play a protective role against oxidative stress-induced damage in epithelial cells (18, 19) Together these results are not only consistent with the crucial role of mitochondrial ATPases in high energy demands of acinar epithelial secretory processes (20), but also suggest dysregulation of ROS-mediated oxidative stress resulting from increased mitochondrial ATPase activity. Resulting ER stress can contribute to functional loss of acinar epithelial cells.

Unresolved ER stress is known to activate NF-*k*B through several mechanisms and leads to inflammatory responses (21). This close interaction between two pathways is also reflected in significantly increased expression of gene encoding NF-*k*B protein (*Nfkb2*) (Figure 2B) in TSP-1^-/-^ acinar epithelial cells consistent with previously reported formation of inflammasomes in these cells and associated increased levels of inflammatory cytokine IL-1β (22). One of the downstream effects of ER stress is the transcription factor ATF6-driven production of ER chaperones, including calreticulin, as part of unfolded protein response (UPR) to enhance the protein folding capacity and mitigate accumulation of misfolded proteins (23, 24). In TSP-1^-/-^ acinar epithelial cells upregulation of both *Atf6* and calreticulin encoding *Calr* gene was detected (Figure 2B). This change was also confirmed by increased detection of calreticulin protein by immunostaining (Figure 2D).

Additionally, in TSP-1^-/-^ acinar epithelial cells we detected a significant downregulation of *Aqp5* gene (Figure 2A), which encodes water channel aquaporin 5 (AQP5) along with a significant downregulation of tight junction proteins like claudins (*Cldn3, Cldn4, Cldn7, Cldn8* and *Cldn34c2*) that are associated with controlled movement of water that is essential for fluid secretion (25-27). These changes reflect disruption of vital components of water transport across the membrane and tear secretion and are consistent with dysregulated AQP5 reported in LG of Sjögren’s patients (28). Furthermore, in TSP-1^-/-^ acinar epithelial cells we noted significant downregulation of *Pigr* gene (Figure 2E) that encodes the polymeric immunoglobulin receptor (pIgR) required for transcytosis of dimeric IgA produced by glandular plasma cells across acinar epithelial cells for delivery as sIgA into tear fluid (29). This finding is further supported by significantly reduced levels of sIgA in tears of TSP-1^-/-^ mice (Figure 2F) as observed in Sjögren’s patients (30). Collectively, these results identify molecular mechanisms that contribute to disrupted secretory function of acinar epithelial cells in TSP-1^-/-^ LGs and the development of autoimmune inflammatory response.

### Disruption of ion transport and calcium signaling contribute to secretory dysfunction of TSP-1 deficient duct epithelium

Duct epithelial cells of the LG participate in tear secretion by fine tuning primary protein-rich lacrimal fluid produced by acinar epithelial cells (6, 31). This modification involves the addition and regulation of water and electrolytes like potassium (K+) and chloride (Cl-) ions, to adjust the composition, osmolarity and volume of final tear fluid. Some of the ion transporters used to achieve this adjustment include Na+/K+ ATP-ase (NKA), Na+/K+/Cl-co-transporter type 1 (NKCC1), cystic fibrosis transmembrane conductance regulator (CFTR) and Epithelial Na+ channel (ENaC). Genes encoding these transporters are *Atp1a1, Atp1b1* (NKAa1 and b1 subunits), *Slc12a2* (NKCC1), *Cftr* (CFTR) and *Scnn1a, Scnn1b* and *Scnn1g* (ENaC a, b and g subunits). Additionally, many other transporters are involved in tear secretion. In our study, we detected significant downregulation of NKA, NKCC1, CFTR, ENaC encoding genes, potassium channels encoded by genes *Kcnj16, Kcnq1*, Sodium bicarbonate transporter *Slc4a11*, Sodium hydrogen exchanger *Slc9a1* and mitochondrial solute carriers that transport ADP/ATP *Slc25a4, Slc25a5, Slc25a16, Slc25a21* (Figure 3A, B, boxed). Significantly reduced expression of ENaC was confirmed by immunostaining as shown in figure 3C. Additionally, we detected downregulated expression of mitochondrial ATPases in TSP-1^-/-^ duct epithelial cells (Figure 3A). Reduced expression of mitochondrial ATPases most notably results from mitochondrial dysfunction and impaired electron transport chain which limits ATP production. This observed gene profile was also supported by a significant downregulation of genes associated with mitochondrial electron transport related pathways (figure 3D). Additionally, significant downregulation of *Camk2n1* that encodes calcium/calmodulin dependent protein kinase II inhibitor was noted in TSP-1^-/-^ duct epithelial cells (figure 3E). Together these results support a significant loss of secretory function of TSP-1^-/-^ duct epithelial cells that likely contributes to compromised tear composition and correlates with ocular surface disease reported in these mice.

**Figure 3:**
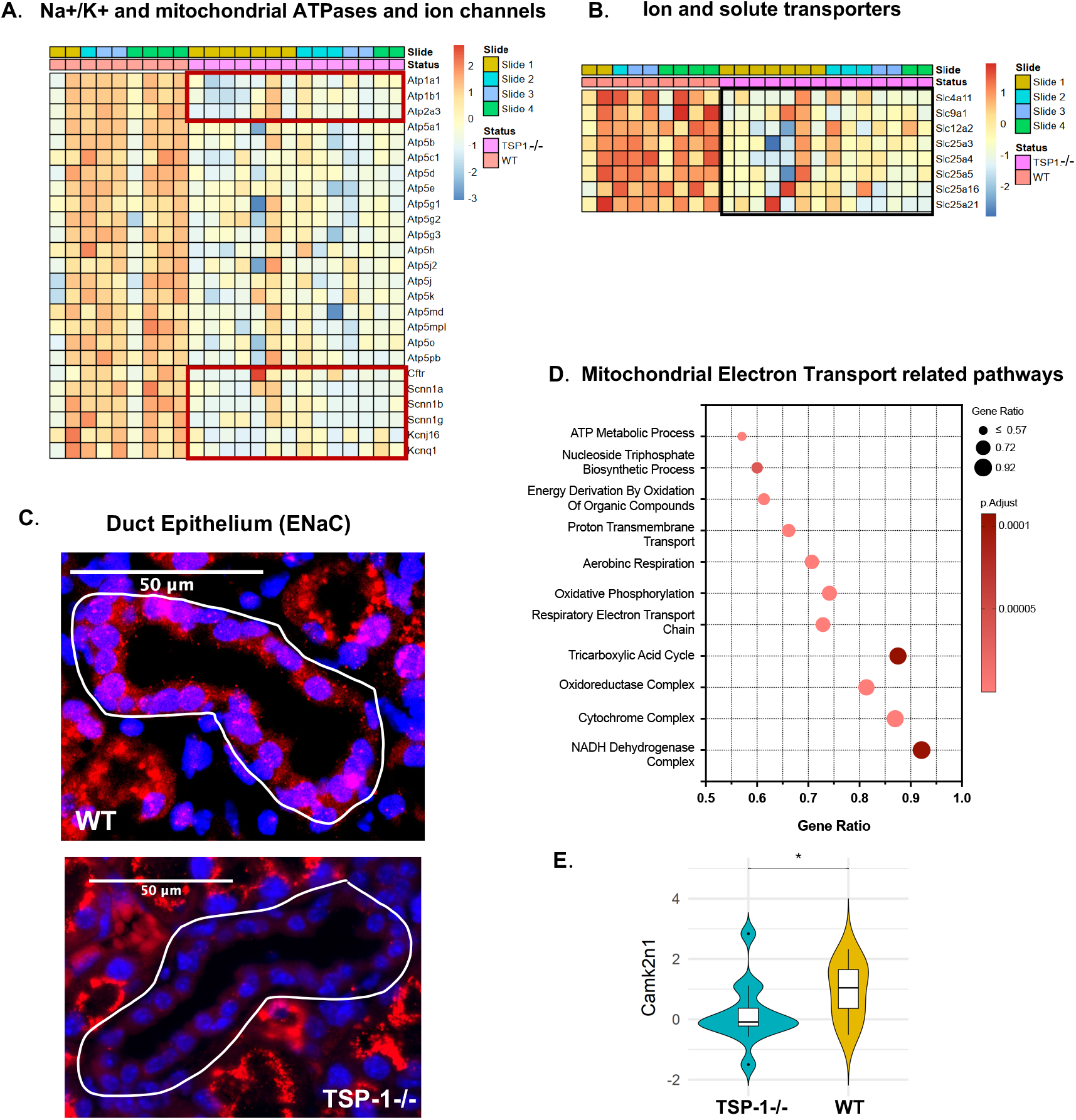
Ion transport and calcium signaling are altered in TSP-1 deficient duct epithelial cells. **(A & B)**. Heatmaps illustrating differential expressions of ion, water, and solute transport-related genes in duct epithelial cells. **(C)** Representative immunofluorescence images of ENaC (epithelial sodium channel) (red) and nuclei (blue) in duct epithelium. White outlines highlight ductal structures. **(D)** Pathways enriched in significantly downregulated DEGs. **(E)** Violin plot comparing *Camk2n1* expression in duct epithelial cells (*q-value <0.05).

### Impaired interactions of TSP-1 deficient myoepithelial cells with acinar epithelial cells underlies structural and functional damage of lacrimal gland

In addition to contractile capacity of MECs that is critical for the propulsion of tear fluid, their regenerative potential (32) is believed to play a significant role in maintenance of the glandular structure (33, 34). Spatial profiling data offered us a unique opportunity to interrogate interactions among MECs and acinar epithelial cells in WT and TSP-1^-/-^ LGs. We identified spatially defined receptor-ligand pairs co-expressed across ROIs in WT and TSP-1^-/-^ glands. As shown in figure 4A, although some receptor-ligand pairs were well correlated in both glands, many pairs were differentially correlated between WT and TSP-1^-/-^ tissues. In normal WT LGs, highly correlated significant interactions included those between MEC-derived laminin-1 and integrin receptors on acinar epithelial cells (e.g. Lamb3 -> Itga6, Lama2 -> Itga6, Lama5 ->Itgb1, Lama5 ->Itgb4, Lama5 -> Dag1) known to promote acinar cell polarity, survival and differentiation as well as interactions between adhesion molecules (Ceacam1 -> Ceacam1, Pcdhgb7 -> Pcdhgb7, Cldn3 - > Cldn3) important for the structural and functional integrity of LGs (35). Significant correlations between growth factors and receptors like Egf ->Egfr and Fgf -> Fgfr2 in WT LGs highlight EGFR signaling important in secretory function of acinar epithelial cells and the role of growth factor FGF in promoting proliferation and migration of MECs (36, 37). Lack of some or relatively fewer such interactions among significantly correlated receptor-ligand pairs in TSP-1^-/-^ tissue coincides with the structural and functional loss observed in TSP-1^-/-^ glands (7). The lack of significant Egf - > Egfr and Cldn3 -> Cldn3 correlations between TSP-1^-/-^ MECs and acinar epithelial cells is supported by significantly reduced expression of *Egf* in TSP-1^-/-^ MECs (1.7-fold, adj. p<0.05) and that of *Cldn3* in TSP-1^-/-^ acinar epithelial cell (1.5-fold, adj. p<0.05). Although we did not detect significantly reduced expression of laminin-1 in TSP-1^-/-^ MECs, significantly increased expression of laminin-degrading metalloproteinase, *Mmp2*, was detected in TSP-1^-/-^ acinar epithelial cells (1.7-fold, adj. p<0.05) that correlates with the pattern of receptor-ligand interactions.

**Figure 4:**
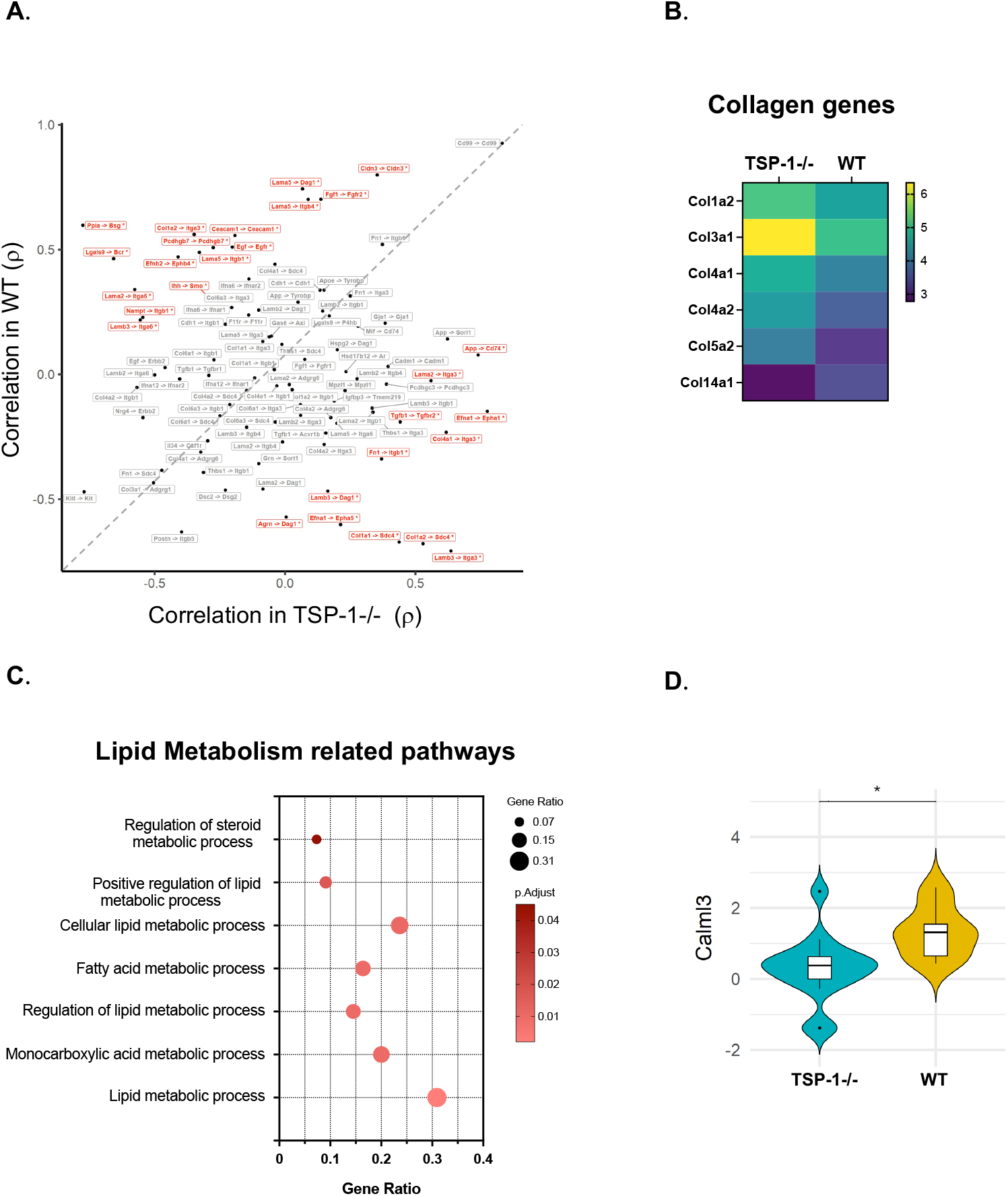
Changes in cellular interactions between MECs and acinar epithelial cells and in MEC contractile function reflecting structural and functional dysfunction in TSP-1 deficient glands. **(A)** Spatially correlated receptor-ligand pairs across Acinar: MEC. Spearmen’s rank correlation coefficients are plotted, and significant interactions are shown in red. **(B)** A heatmap showing the significantly altered expression of various collagen genes. **(C)** Pathways enriched in downregulated DEGs **(D)** A violin plot comparing *Calm3* expression (*q-value <0.05).

Unlike laminin-1 in WT glands, in TSP-1^-/-^ glands highly correlated ECM proteins bound by integrins predominantly included fibronectin and collagens that are associated with epithelial repair following injury (38-40). In fact, significantly increased receptor-ligand correlations (e.g. Tgfb1 -> Tgfbr2, Fn1 ->Itgb1, Col4a1 ->Itga3, Col1a2 -> Sdc4) in TSP-1^-/-^ gland suggestive of profibrotic process is accompanied by increased expression of collagens (figure 4B) known to play distinct roles in ECM remodeling during tissue repair (41). However, expression of type XIV collagen, *Col14a1*, known to regulate assembly and organization of collagen fibrils (42, 43) required to create ECM scaffold, was significantly reduced indicating impaired ECM organization. Furthermore, significant correlation between Amyloid precursor protein and MHC invariant chain (App -> Cd74) is consistent with an increased CD74 expression in TSP-1^-/-^ acinar epithelial cells (1.7-fold, adj. p<0.05). Similar increased epithelial CD74 expression is reported in diverse tissue injury diseases and inflammatory responses including autoimmune diseases (44-46).

We also detected enrichment of pathways associated with lipid metabolism among differentially downregulated genes in TSP-1^-/-^ MECs (figure 4C). As lipid metabolism is known to support smooth muscle contraction through various mechanisms (47, 48) this finding suggests a potential loss of contractile responses in TSP-1^-/-^ MECs. This possibility is also supported by significantly reduced expression of calcium sensor encoding gene, *Calml3* (figure 4D) further corroborating previously reported *in vitro* observations of altered calcium signaling and contractile responses in primary cultures of TSP-1^-/-^ MECs (49). Collectively our findings indicate that disruption of distinct interactions of MECs with acinar epithelial cells and their contractile function significantly contribute to the structural and functional loss observed in TSP-1^-/-^ LGs.

### Periductal immune infiltrates in TSP-1 deficient lacrimal glands form GCs

As reported previously we observed periductal and perivascular lymphoid aggregates characteristic of SjD pathology in TSP-1^-/-^, but not WT LGs. In our DSP study CD45 staining in ROIs containing immune infiltrates was very weak which we confirmed by staining additional slides (supplementary figure 3). Among immune cells low CD45 expression is reported in B cell subsets that is related to their enhanced survival and autoimmunity (50-52). Staining of these aggregates with B cell marker (B220) confirmed B cell predominance (figure 5A). Expression of markers associated with germinal center B cells (GC B) and T follicular helper (Tfh) cells was detectable in the transcriptomic profile of such aggregates (figure 5B) suggesting the presence of activated B cells. Moreover, pathway analysis of DEGs in the aggregate revealed significant enrichment of pathways associated with the formation of germinal center (figure 5C) suggesting lymphoid aggregates to be active germinal centers (GCs). We also detected CD4 positive cells that stained for Bcl6, the lineage-defining transcription factor of Tfh cells. These cells were found in GC as well as among periductal immune cells in TSP-1^-/-^ LGs (figure 5D) further validating our gene expression profile of lymphoid aggregates. To determine if the observed GCs in TSP-1^-/-^ LGs correlate with spontaneous splenic GC formation that is commonly associated with systemic autoimmunity, we evaluated WT and TSP-1^-/-^ spleen sections. Relative to WT spleen, more activated follicles marked by the presence of B220+Bcl6+ (yellow) GC B cells and B220-Bcl6+ (green) Tfh cells are seen in TSP-1^-/-^ spleen (figure 5 E-a-f). We further confirmed Tfh cells as CXCR5+CD4+ (yellow) cells as seen in figure 5E panels g-j near the periphery and within the follicle in TSP-1^-/-^ spleens, in comparison to WT spleen. Together these results support the presence of ectopic GCs in LGs that correlates with spontaneous splenic GCs in TSP-1^-/-^ mice consistent with their systemic autoimmunity.

**Figure 5:**
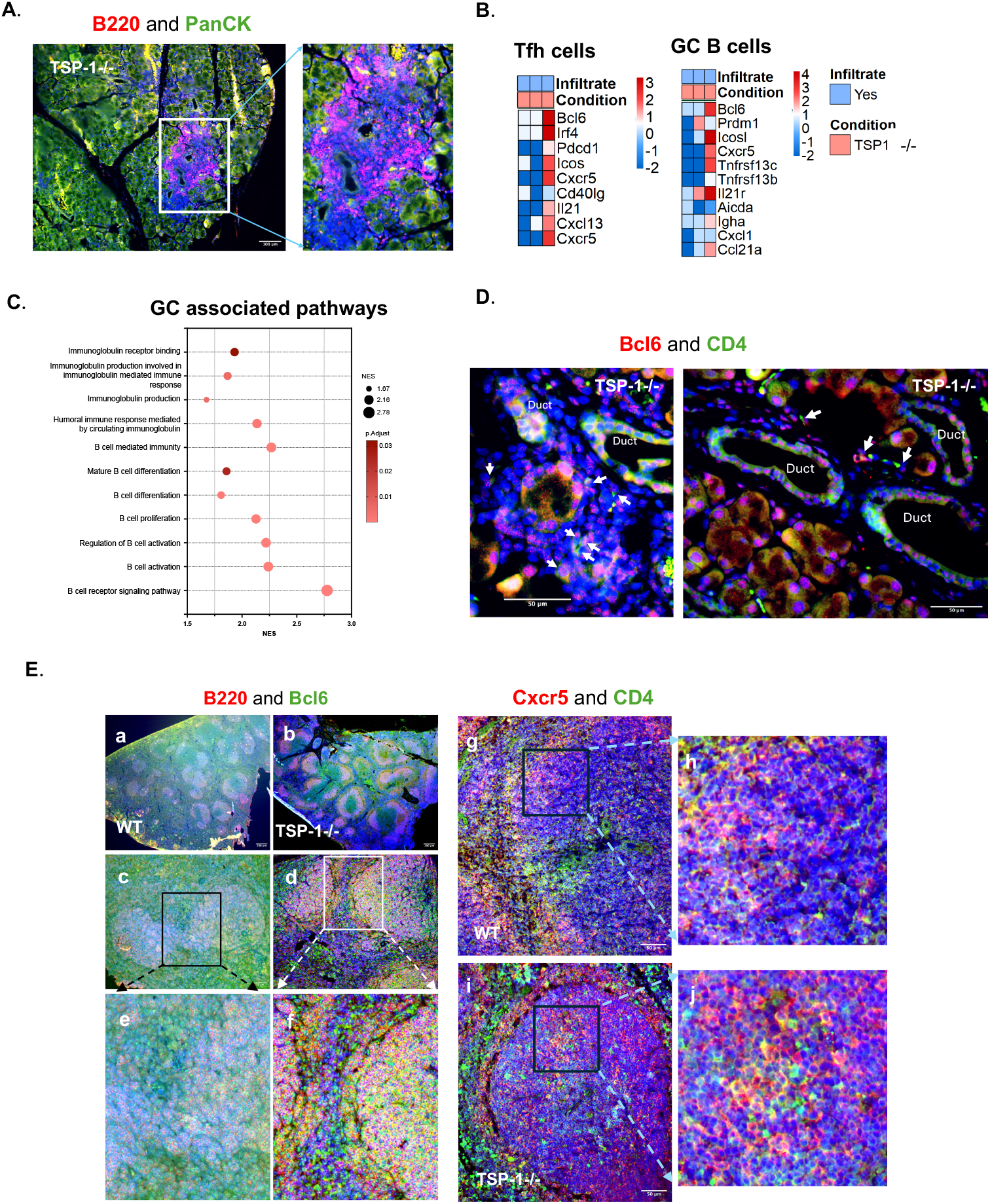
Spontaneous germinal centers containing B cells and Tfh cells in the lacrimal gland and spleen of TSP-1 deficient mice. **(A**) Immunofluorescence staining for B220 and PanCK showing predominance of B cells in periductal infiltrates detected in TSP-1^-/-^LG; **(B)** Heatmaps of gene expression profiles of GC B and Tfh cells in ROIs containing infiltrates; **(C)** Pathways enriched in significantly upregulated DEGs in periductal immune cells; **(D)** Representative immunofluorescence images showing Tfh (Bcl6+CD4+) cells in the periductal infiltrates (white arrows); **(E)** Immunofluorescent images of spleen sections stained for GC B (Bcl6+B220+) cells (subpanels a-f: boxed regions in c & d enlarged in e & f respectively below to show GCs in the follicle and Tfh cells showing only Bcl6 staining at the periphery and within the follicle) and Tfh (CXCR5+ CD4+) cells (subpanels g-i: boxed regions in g & i are enlarged in h & j respectively on the right to show Tfh cells in the follicle). Co-expression of markers indicated by yellow color.

### Immune cells adjacent to duct epithelial cells in TSP-1 deficient lacrimal gland include antigen presenting cells capable of recruiting Tfh and GC B cells

In our DSP study ROIs containing ducts included CD45 staining in the proximity of PanCK stained duct epithelial cells in both WT and TSP-1^-/-^ LGs as shown in figure 6A. There were some duct ROIs with higher CD45 staining intensity than others. Relatively more duct ROIs had higher CD45 staining intensity in TSP-1^-/-^ glands (40%) than in WT glands (22%). We compared transcriptional profile of duct ROIs with high and low CD45 staining intensities. The pattern of genes associated with antigen presenting cells (APCs) correlated with the CD45 staining intensity in duct ROIs (figure 6B). Such a correlation suggested a presence of APCs among immune cell population residing in spaces between duct epithelial cells revealing as-yet-unknown spatial relationship between duct epithelial cells and APCs in LGs. There is evidence supporting APCs (either monocyte/macrophages or DCs) as a source of CXCL13 during inflammation due to their heightened sensitivity to inflammatory stimuli through Toll-like receptor (TLR) pathways including TLR7/9 and TLR4 and cytokines like TNF-α (53, 54). Considering the periductal nature of infiltrates that include CXCR5-expressing Tfh cells as well as GC B cells in TSP-1^-/-^ LGs, we compared expression of chemokine, CXCL13, and inflammatory molecules associated with its stimulation in immune cells located near duct epithelial cells in TSP-1-/- and WT LGs. As shown in figure 6C, in several ROIs in TSP-1^-/-^ LGs, increased expression of selected genes was detectable, and immunostaining confirmed CXCL13 co-localization with CD45 (figure 6D) thus supporting the development of observed periductal GCs. Furthermore, we also detected significantly increased expression of MHC class II molecules in duct epithelial cells in TSP-1^-/-^ LGs as compared to WT LGs (figure 6E) suggesting their potential of presenting autoantigens during glandular inflammation.

**Figure 6:**
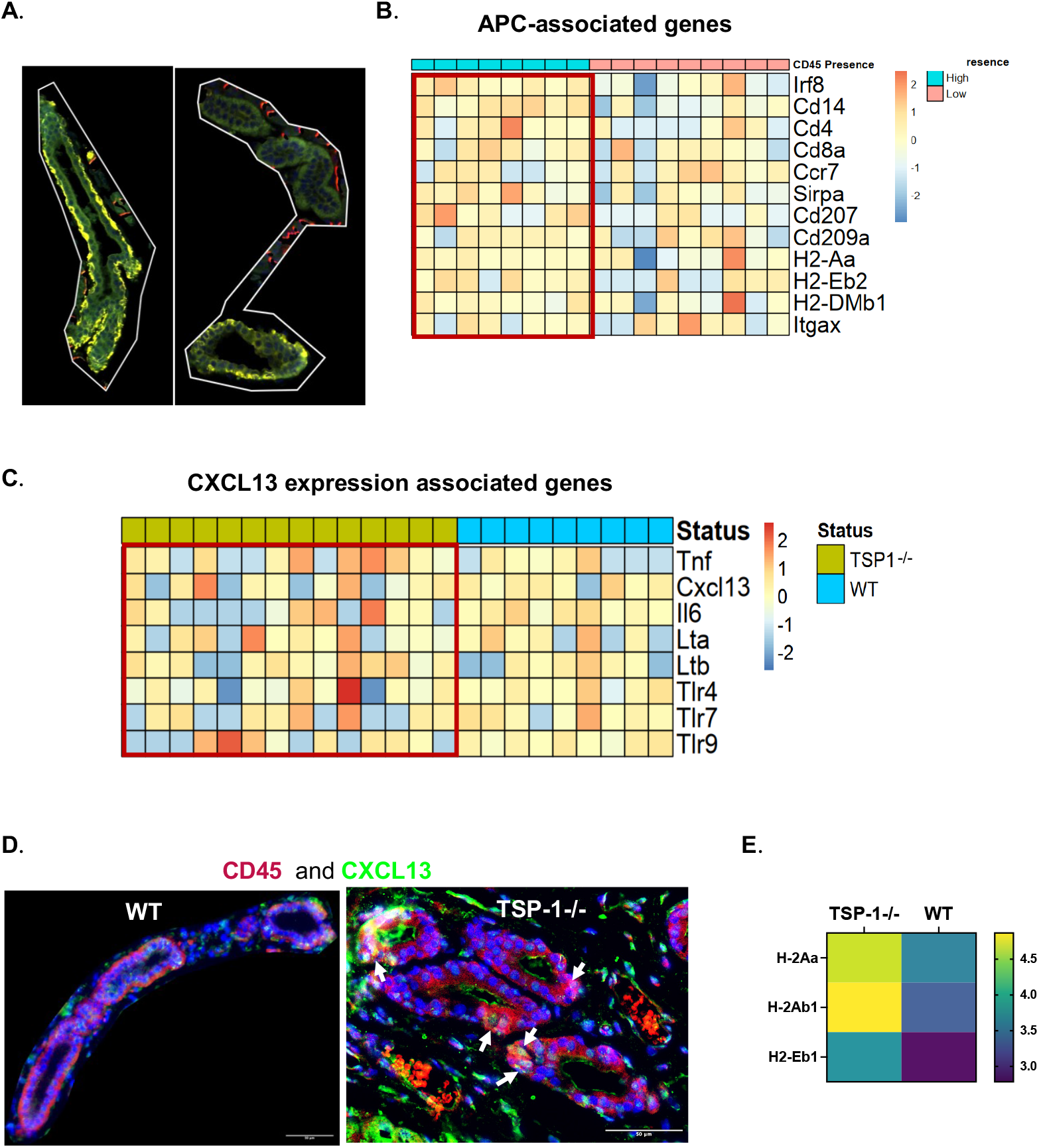
Spatial relationship of duct epithelial cells with antigen-presenting cells and their pro-inflammatory gene signatures in TSP1 deficient lacrimal glands. **(A)** Representative images showing ROIs containing PanCK (green) positive duct epithelial cells and CD45 (yellow) positive immune cells with variable (high and low) staining intensities; **(B)** Heatmap of APC-associated gene expression in ROIs containing ducts with high vs. low CD45 staining intensity; **(C)** Heatmap analysis of all ROIs containing ducts for genes associated with CXCL13 induction in APCs; **(D)** Representative images of WT and TSP-1^-/-^ ducts showing CXCL13 co-localization with CD45 (white arrows) in the latter; **(E)** Heatmap showing significantly altered expression of MHC class II genes in duct epithelial cells.

### Duct epithelial cells in TSP-1 deficient lacrimal glands provide the microenvironment to generate GCs

The proximity of duct epithelial cells to APCs suggested a possibility of duct epithelial cell-derived factors shaping the local immune response induced by APCs. In autoimmune pathology, IL-6 is a key cytokine known to drive polyclonal activation of B cells to facilitate emergence of autoreactive clones, promote differentiation of Tfh cells that help in GC formation and act synergistically with CXCL13 to recruit and retain B and Tfh cells (55, 56). We detected several duct epithelial ROIs in TSP-1^-/-^ LGs with relatively higher expression of IL-6. To validate this finding and determine if duct epithelial cells contribute to microenvironment supportive of periductal GC formation, we generated primary cultures of LG-derived duct epithelial cells as described in methods. These cultures were validated by confirming their expression of epithelial cell (PanCK) and duct epithelial cell (CFTR) markers (57) and the absence of myoepithelial and acinar cells as evident from minimal to no detectable immunostaining for α-SMA and Rab3d respectively (supplementary figure 4). Duct epithelial cell cultures derived from WT and TSP-1^-/-^ LGs were used to collect 24 h culture supernatants to compare their secretion of IL-6. As shown in figure 7A, significantly increased IL-6 levels were detectable in TSP-1^-/-^ duct epithelial cell cultures as compared to WT control cultures. Another way epithelial cells are known to influence the local immune response is by functioning as non-professional APCs by expressing MHC class II molecules in response to IFN-γ and participate in activation of effector CD4+ T cells (58). We next assessed if TSP-1^-/-^ duct epithelial cells differed from WT controls in their ability to respond to IFN-γ by determining their expression of MHC class II by RT-PCR after exposure to IFN-γ as described in methods. As shown in figure 7B, significantly increased expression of MHC class II was detectable in TSP-1^-/-^ duct epithelial cells as compared to the WT controls. Collectively these results support the potential of duct epithelial cells to shape local immune response by facilitating the formation of periductal GCs in TSP-1^-/-^ LGs.

**Figure 7:**
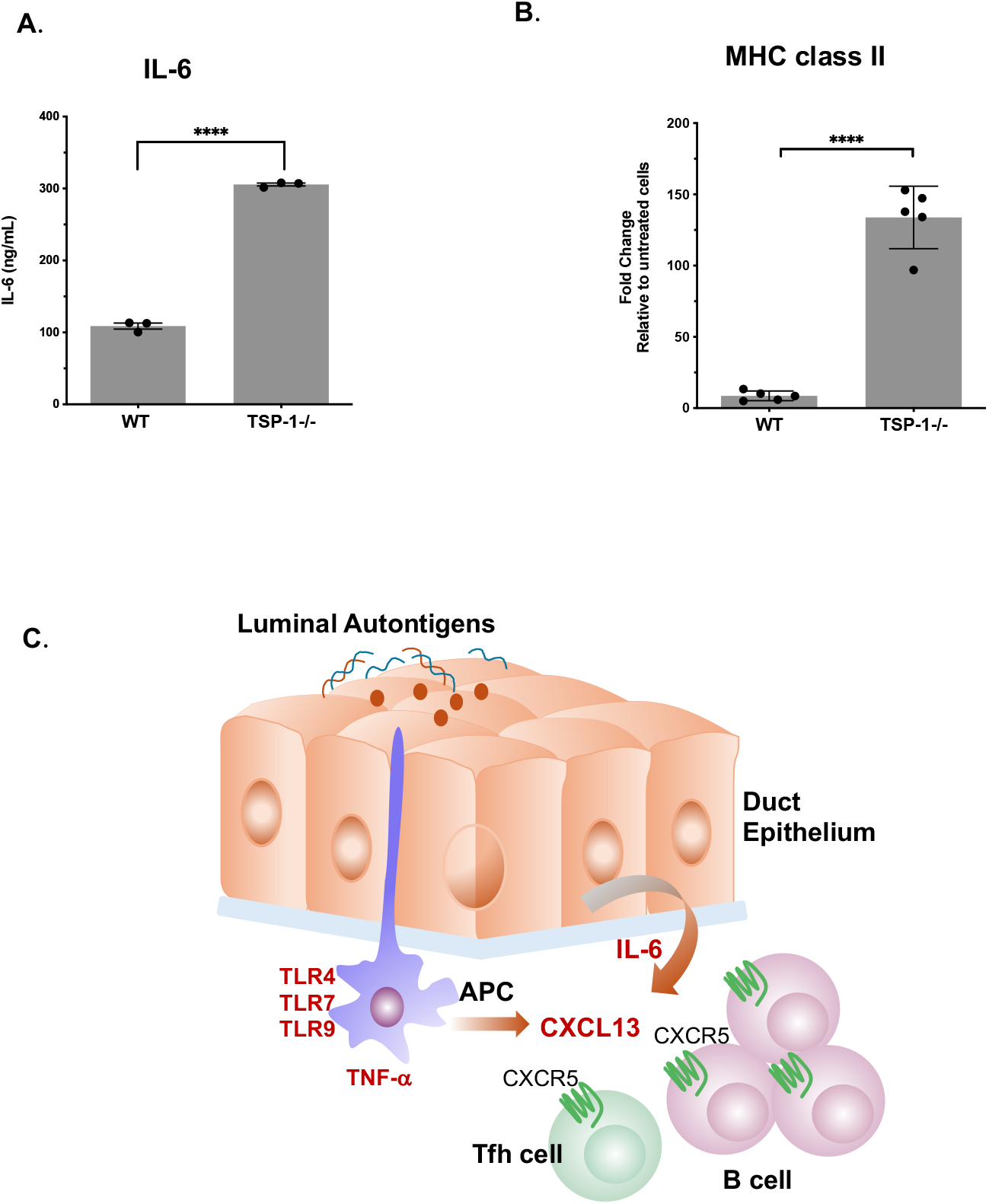
Duct epithelial cells from TSP-1 deficient lacrimal gland secrete increased IL-6 and express higher levels of MHC class II under inflammatory conditions. **(A)** Levels of IL-6 detected in 24 h supernatants of primary cultures of duct epithelial cells; **(B)** Expression of MHC class II gene in IFN-g-stimulated primary duct epithelial cell cultures; Data presented as mean ± SEM, ****p<0.001; **(C)** Schematic representation of molecular mechanisms underlying the development of periductal immune infiltrates and spontaneous GC formation. Capturing luminal autoantigens by APCs adjacent to duct epithelial cells can induce their expression of pro-inflammatory molecules TNF-a and CXCL13 to help recruit CXCR5-expressing Tfh and GC B cells. Duct epithelial cell-derived IL-6 can provide microenvironment permissive for the proliferation of recruited cells and facilitate formation of spontaneous periductal GC.

## Discussion

In this study we used DSP of LG tissues derived from healthy and SjD mouse model to spatially define interactions between epithelial cells and immune cells that contribute to autoimmune response and glandular pathology. Our approach not only provided *in situ* cell-type representation comparable to that reported with scRNA-seq method but additionally helped detect critical spatially relevant molecular mechanisms underlying structural and functional loss of LGs in SjD. Furthermore, this study identifies molecular mechanisms supportive of a tissue microenvironment that facilitates the development of periductal infiltrates and germinal centers in LG that characterize SjD pathology.

Endoplasmic reticulum (ER) stress is increasingly recognized as a key factor in the development and progression of autoimmune diseases including rheumatic diseases (59). Unfolded protein response (UPR), activated by ER stress, plays an essential role in regulation and maintenance of innate as well as adaptive immunity. However, activation of ER stress in LGs with Sjögren’s pathology was not clear until now. Selective examination of transcription profile of acinar epithelial cells in our study enabled detection of upregulated ER stress pathway in TSP-1^-/-^ LGs similar to those reported in salivary glands of patients with primary SjD (60) and consistent with an observed SjD-like phenotype in mice with ablated ER-resident chaperone protein (61). Furthermore, ER stress can activate pro-inflammatory NF-*k*B including NLRP3 inflammasome and related cell death (21, 62). Our findings are corroborated by previously observed formation of inflammasomes in TSP-1^-/-^ LG epithelial cells (22) and their increased apoptotic cell death (7).

Additionally, the increased expression of calreticulin in acinar epithelia of TSP-1^-/-^ glands is one of the downstream effects of their ER stress (23). In particular, calreticulin expression is associated with immunogenic cell death (ICD) that leads to aberrant immune activation and autoantibody production to exacerbate autoimmunity (63) as observed in TSP-1^-/-^ mice. In fact, calreticulin is reported to participate in antigen presentation of the Ro60 (SSA) epitopes (64, 65). Therefore, our current findings not only explain the detection of anti-Ro antibodies in sera of TSP-1^-/-^ mice (7), but are also congruent with the detection of anti-calreticulin autoantibodies in the sera of SjD patients (66). Furthermore, the structural damage to LGs in SjD which is also reflected in reduced expression of *Pigr* (cell surface receptor for IgA transcytosis) and validated by the disrupted delivery of sIgA into tears. There is growing evidence implicating sIgA regulation of commensal microbiota in mucosal inflammatory disease (67, 68). Reduced tear sIgA in TSP-1^-/-^ mice is consistent with their altered ocular commensal microbiota as reported previously (69) which is also implicated in SjD pathogenesis. Additionally, our results suggest tear sIgA may serve as a useful biomarker that reflects glandular epithelial damage.

In addition to acinar epithelial cells, our study identified molecular mechanisms underlying functional loss of duct epithelial cells that are critical for maintaining balanced tear composition that maintains integrity of the ocular surface. Duct epithelial cells modify tear fluid by regulating ion and water components via various ion transporters/channels to produce K^+^ and Cl^-^ rich tears (31). In a rabbit model of SjD, reduced expression of Na^+^/K^+^ ATPases, Chloride channel and epithelial Na^+^ channel in duct epithelial cells correlated with their altered LG secretion and ocular surface disease (70, 71). Similarly, our study detected reduced expression of these channels in duct epithelial cells which correlates LG functional loss and ocular surface disease in TSP-1^-/-^ mice. Furthermore, reduced expression of mitochondrial ATP synthases and electron transport chain related pathways in duct epithelial cells are suggestive of mitochondrial dysfunction, which has been implicated in autoimmune disease pathogenesis (72). Malfunctioning mitochondria can serve as a source of autoantigens in the form of mitochondrial-nucleic acids and release mitochondrial danger associated molecular patterns (DAMPs) that can contribute to and intensify inflammatory responses (73, 74). Also, reduced expression of calcium signaling related molecule (*Calml3*) in TSP-1^-/-^ MECs and acinar epithelial cells supports their functional loss consistent with previously reported loss of contractile and secretory responses respectively (7, 49, 75). Uniquely, our study also detected profibrotic changes in cellular interactions between MECs and acinar epithelia cells in TSP-1^-/-^ LGs along with altered interactions of ECMs suggesting dysregulated ECM organization that reflects loss of structural integrity in LG further confirming their overall functional loss. These mechanisms may represent early markers in progressive fibrotic changes that are correlated with dry eye in patients as well as glandular atrophy noted in advanced SjD (76, 77).

Chronic LG inflammation in SjD is marked by characteristic periductal and perivascular infiltrates containing B and T cells that often form foci sometimes organizing into GCs that facilitate local B cell activation to perpetuate autoimmunity within the gland (14, 78). Locally produced autoantibodies contribute to tissue damage and inflammation. Molecular and cellular cues that sustain periductal immune infiltrates or GCs in LGs are not completely defined. Particularly contribution of epithelial cells to microenvironment that fosters formation of GCs is not known. In our study we detected predominance of B cells in periductal infiltrates in TSP-1^-/-^ LGs like those reported in SjD patients and other mouse models (14, 79, 80). The presence of spontaneously formed GCs (containing GC B and Tfh cells) in the LG and spleen of TSP-1^-/-^ mice is also consistent with the presence of Sjögren’s specific autoantibodies in this SjD model (7). Our spatial profiling approach also enabled detection of APCs in the proximity of duct epithelial cells. To the best of our knowledge this is the first study to reveal such a location of APCs in LGs supporting their potential to sample autoantigens released in the lacrimal fluid (figure 7D). Expression of signaling molecules TLRs and TNF-α known induce Tfh and GC B cell-attracting chemokine, CXCL13, by some of these APCs also helps explain the periductal location of infiltrates and GCs in LGs (53, 54). Additionally, we report for the first time the ability of duct epithelial cells in LG to express MHC class II suggesting their participation in immune surveillance and shaping of the local immune response. Such epithelial MHC class II expression is reported in mucosal tissue in response to IFN-γ (58, 81). Our *in vitro* experiments confirmed similar responsiveness of TSP-1^-/-^ duct epithelial cells. Furthermore, their ability to produce increased levels of IL-6 explains their ability to foster microenvironment that sustains autoreactive B cell responses indirectly by facilitating Tfh differentiation to support GC reactions and directly by promoting B cell differentiation into antibody secreting plasma cells (56, 82). Collectively our findings elucidate mechanisms underlying the development of periductal immune infiltrates in LG pathology seen in SjD.

In summary, spatial transcriptomic profiling approach successfully identified previously reported epithelial cell subtypes and further helped us expand on prior studies by dissecting cellular ecosystem in LGs under both normal and disease condition. In addition to detecting mechanisms underlying functional loss of LG epithelial cells our study identifies (1) their active contribution in promoting and sustaining autoimmune inflammation beyond serving as passive targets, (2) spatially relevant molecular drivers of GC formation, (3) potential early markers of profibrotic changes, (4) potential tear biomarker indicative of glandular damage.

## Materials and Methods

### Animals

Twelve-week-old male C57BL/6 and B6.129S2-Thbs1<tmlHyn>/J (TSP1^-/-^) mice were purchased from Jackson Laboratories (Bar Harbor, ME, USA). The Institutional Animal Care and Use Committee (IACUC) at Boston University School of Medicine, Boston, approved the animal studies described in this manuscript (Protocol number IPROTO202200000063) in accordance with the National Institutes of Health (NIH) guide for the care and use of laboratory animals. All animal experiments were conducted in accordance with the Association for Research in Vision and Ophthalmology (ARVO) statement for the Use of Animals in Ophthalmic and Vision Research.

### Digital Spatial Profiling (DSP)

Serial sections of formalin-fixed paraffin-embedded (FFPE) LG tissues harvested from WT (n=3) and TSP^-/-^ (n=4) mice were cut (5 µm thick) and transferred to slides. One set of slides was stained for H&E and another set of slides with consecutive sections were submitted for spatial profiling to the Beth Israel Deaconess Medical Center (BIDMC) Spatial Technologies Unit (STU), MA, USA. The GeoMX Digital Spatial Profiler (DSP) platform was used to generate spatially resolved transcriptomic data, specifically the Whole Transcriptome Atlas (WTA) mouse RNA probe set for Illumina Systems (GMX-RNA-NGS-MsWTA-4) assay. This assay includes RNA probe set for 21,000+ transcripts for mouse protein coding genes plus negative controls designed for Illumina NGS readout with the Seq Code library prep. To visualize tissue structural components and microenvironment, a set of fluorophore-conjugated morphology markers were used. These included Alexa Fluor 532 (green)-conjugated anti-pan-CK (AE1+AE3, Novus, Cat.# NBP2-33200), Alexa Fluor 594 (yellow)-conjugated anti-CD45 (EM-05, Novus, Cat.# NBP1-44763AF594) and Alexa Fluor 647 (red)-conjugated anti-Smooth Muscle Actin (SP171, Abcam, Cat.# ab267537) and nuclear stain SYTO13 (blue). These markers were used to delineate the nuclear, epithelial (acinar and duct), immune and myoepithelial compartments. Immunofluorescence imaging, region of interest (ROI) selection, segmentation into marker-specific areas of interest (AOI) and spatially indexed barcode cleavage and collection were performed on a GeoMx Digital Spatial Profiling instrument (Nanostring). Selected ROIs were exposed to ultraviolet (UV) light to cleave DNA tags in a region-specific manner. The released indexing oligos were then collected using microcapillary aspiration and dispensed into a microplate as per segmentation of each ROI.

Four consecutive sections per tissue (total 16 of TSP-1^-/-^ and 12 of WT) were assessed. Approximately, 6-10 ROIs and 11-14 AOIs per tissue sample (total 34 ROIs and 57 AOIs for TSP-1^-/-^ and 21 ROIs and 34 AOIs for WT) were collected. Library preparation was performed as per manufacturer’s instructions which included PCR amplification to add Illumina adapter sequences and unique dual sample indices. Sequencing was performed to a minimum of 350M PE reads by pooling all 91 uniquely indexed WTA AOIs and sequencing on Illumina NovaSeq 6000 S4 flowcell in a 2x150bp PE, dual index, 8bp i5 and i7 index length configuration.

### DSP Data preprocessing and analysis

Processing of RNA sequencing files (FASTQ files) from Illumina sequencer according to parameters defined in the configuration file generated from the GeoMX DSP run was performed using the GeoMx NGS Pipeline v2.0 (Bruker Spatial Biology, formerly Nanostring Technologies, Inc., WA, USA) to produce digital count conversion (.dcc) files. Counts were loaded into R Statistical Software (v4.3.2) (83), whereupon they were normalized and evaluated for differential expression using DESeq2 (84), filtering out all unannotated genes (Rik and Gm), applying a median count threshold of 5 per group for independent expression filtering, and utilizing Storey’s q-values for multiple hypothesis correction (85). Subsequently ClusterProfiler (86) was used to carry out over-representation (ORA) and gene-set enrichment (GEO) analysis for pathways. Gene visualizations were done using variance stabilized transform normalized counts or log-counts. Cell type proportions in ROIs were estimated using Multi-subject Single Cell (MuSiC) deconvolution analysis toolkit (87) with a publicly available dataset GSE132420 as a reference (14). For receptor-ligand correlation across ROIs, known receptor-ligand pairs were obtained from CellChatDB (88) and potential receptor-ligand pairs were quantified using the Spearman rank correlation between paired segments within each ROI. To compare magnitude of correlations between WT and TSP-1-/-tissues differential correlation analysis of cell types across conditions was performed and cell type rich segments were paired per ROI per condition. The cocor R package (89) was then used for statistical comparison of correlations. Plots were generated using ggplot2 (90), ggpubr (91), and pheatmap (92).

### Immunofluorescence staining

To validate the expression of selected genes of interest, 5µm thick FFPE sections of LGs were immunostained using specific antibodies. For these slides with sections were baked at 60^0^C overnight, deparaffinized with xylene (10min; twice) and rehydrated and subjected to Heat-induced antigen retrieval (HIER) in citrate-based buffer pH 6 (Vector laboratories, California, USA) for 25 minutes in a microwave. After antigen retrieval slides were photobleached and sections were blocked with 10% normal goat or donkey serum for 1 hour followed by 2% bovine serum albumin (Sigma Aldrich, St Louis, USA) in 0.3% Triton-X100 in PBS for 30 minutes at room temperature (RT). Sections were incubated overnight at 4^0^C with 1:50 dilution in PBS-BSA of primary antibodies listed in supplementary table 1, followed by three washes with PBS-0.05%Tween20 (2 min each). Tissues were then incubated for 1.5 hour at RT in the dark with fluorescence-conjugated secondary antibody (1:500-1:1000) listed in supplementary table 1, washed three times with PBS-0.05%Tween20 (5 min each) and counterstained with DAPI and mounted using Pro-long gold Antifade Mountant (Invitrogen, USA). For dual-immuno-staining sequential staining was performed (Novus Biologicals Ref). For immunostaining of spleen sections similar steps were followed except in case of CD4 staining Zinc fixative (BD Biosciences, CA, USA) was used without antigen retrieval steps.

In case of primary cell cultures, cells grown on coverslips were fixed in ice-cold methanol for intracellular targets like cytokeratin and smooth muscle actin and with 2% PFA for cell surface target like CFTR. Fixed cells were incubated at RT for 15 minutes with blocking buffer containing 2% BSA/1% Triton X-100 in PBS for intracellular targets and for 1 hour with 2% BSA in PBS for cell surface target. After three washes with PBS, cells were incubated with primary antibodies at RT for 3 hours followed by 1 hour with fluorescence-conjugated secondary antibody. After final washes with PBS cells were mounted in mounting medium containing nuclear stain DAPI for microscopic examination.

Images were acquired using a fluorescence microscope (Nikon Eclipse E800, Nikon, Japan) equipped with a MicroPublisher 6 camera and further processed using Fiji ImageJ software (National Institutes of Health, USA).

### Primary lacrimal gland duct epithelial cell culture

Keratinocyte basal medium (KBM) and RPMI-1640 (Lonza) supplemented with 10% heat inactivated fetal calf serum (Thermo Fisher Scientific, Waltham, MA), 1 mM Sodium Pyruvate, 10 mM HEPES, 100 mg/ml Penicillin-Streptomycin and 0.1 mM NEAA (Sigma Aldrich, St. Louis, MO) were used in 1:1 proportion for cell cultures. Lacrimal gland tissues were minced into small pieces and that were anchored onto scored 48-well culture plates. Three pieces of tissue were anchored per culture well with approximately 75 ml medium to cover the bottom of the well. Culture plates were incubated under routine culture conditions of 5% CO2 at 37^0^C. Medium was replaced every 2-3 days and after tissue pieces adhered to the plastic, medium volume was increased to 100 ml. Cell growth was monitored routinely until it reached >75% confluence in 10-14 days after which explants were removed. Cells were used either for immunostaining or culture supernatants were collected for ELISA. In some experiments, cells were treated for 24 hours with recombinant mouse IFN-γ (10 ng/ml, R&D Systems, Minneapolis, MN).

### RT-PCR

Total RNA isolated from untreated or IFN-γ-treated epithelial cell cultures using TRIzol reagent (Life Technologies, Carlsbad, CA) was used to synthesize cDNA using SuperScript VILO cDNA synthesis kit (Thermofisher Scientific, Waltham, USA) according to the manufacturer’s instructions. For amplification of MHC class II-specific gene transcripts primer set (F-5’-AGG GCA TTT CGT GTA CCA GTT and R-5’-GTA CTC CTC CCG GTT GTA GAT GTA) was used along with primers for the reference gene glyceraldehyde-3-phosphate dehydrogenase primers (F-5’-CGAGAATGGGAAGCTTGTCA-3’ and R-5’-AGACACCAGTAGACTCCACGACAT-3’). Real-time PCR assay was performed using SYBR Green PCR Master Mix (Thermofisher Scientific, Waltham, USA) with amplification reactions set up in quadruplicates and thermal profile: 95°C for three minutes, 40 cycles at 95°C for ten seconds, 54.5°C for ten seconds and 72°C for thirty seconds. Specificity of the amplification reaction was verified by performing melting curve analysis. The threshold cycle values were used to determine quantification of gene expression relative to reference gene GAPDH.

### ELISA

Culture supernatants collected from primary cultures of WT and TSP-1^-/-^ LG epithelial cells were analyzed for IL-6 content using ELISA (eBioscience, CA, USA) according to manufacturer’s instructions. Pilocarpine-induced tear samples collected from 24-wk-old mice were analyzed for their sIgA content using ELISA (Novus Biologicals, CO, USA).

## Statistical Analysis

Student’s unpaired t-test was used to determine significant differences between mean values of experimental and control groups and p<0.05 was considered significant. Statistical analysis was performed using GraphPad Prism 10 software.

## Acknowledgments

This study was supported in part by research grant from NIH/NEI (R56EY035213).

## Notes

### Competing Interest Statement

The authors have declared no competing interest.

## References

1. O. D. Argyropoulou, A. G. Tzioufas, Update on Sjogren’s Syndrome 2018. Mediterr J Rheumatol 29, 193–198 (2018).

2. P. Brito-Zeron et al., Sjogren syndrome. Nat Rev Dis Primers 2, 16047 (2016).

3. J. P. Whitcher et al., A simplified quantitative method for assessing keratoconjunctivitis sicca from the Sjogren’s Syndrome International Registry. Am J Ophthalmol 149, 405–415 (2010).

4. Y. Liang, Z. Yang, B. Qin, R. Zhong, Primary Sjogren’s syndrome and malignancy risk: a systematic review and meta-analysis. Ann Rheum Dis 73, 1151–1156 (2014).

5. M. Ramos-Casals et al., EULAR recommendations for the management of Sjogren’s syndrome with topical and systemic therapies. Ann Rheum Dis 79, 3–18 (2020).

6. B. Walcott, The Lacrimal Gland and Its Veil of Tears. News Physiol Sci 13, 97–103 (1998).

7. B. Turpie et al., Sjogren’s syndrome-like ocular surface disease in thrombospondin-1 deficient mice. Am J Pathol 175, 1136–1147 (2009).

8. L. Contreras-Ruiz, B. Regenfuss, F. A. Mir, J. Kearns, S. Masli, Conjunctival inflammation in thrombospondin-1 deficient mouse model of Sjogren’s syndrome. PLoS One 8, e75937 (2013).

9. R. J. Chaparro et al., Nonobese diabetic mice express aspects of both type 1 and type 2 diabetes. Proc Natl Acad Sci U S A 103, 12475–12480 (2006).

10. E. Clough, T. Barrett, The Gene Expression Omnibus Database. Methods Mol Biol 1418, 93–110 (2016).

11. A. Oyelakin et al., Transcriptomic and Network Analysis of Minor Salivary Glands of Patients With Primary Sjogren’s Syndrome. Front Immunol 11, 606268 (2020).

12. T. Greenwell-Wild et al., Chitinases in the salivary glands and circulation of patients with Sjogren’s syndrome: macrophage harbingers of disease severity. Arthritis Rheum 63, 3103–3115 (2011).

13. Q. Fan et al., Exploring Immune Cell Diversity in the Lacrimal Glands of Healthy Mice: A Single-Cell RNA-Sequencing Atlas. Int J Mol Sci 25 (2024).

14. A. Rattner, J. S. Heng, B. L. Winer, L. A. Goff, J. Nathans, Normal and Sjogren’s syndrome models of the murine lacrimal gland studied at single-cell resolution. Proc Natl Acad Sci U S A 120, e2311983120 (2023).

15. V. Delcroix et al., The First Transcriptomic Atlas of the Adult Lacrimal Gland Reveals Epithelial Complexity and Identifies Novel Progenitor Cells in Mice. Cells 12 (2023).

16. O. E. Ospina et al., Differential gene expression analysis of spatial transcriptomic experiments using spatial mixed models. Sci Rep 14, 10967 (2024).

17. S. L. Goldman et al., The Impact of Heterogeneity on Single-Cell Sequencing. Front Genet 10, 8 (2019).

18. Y. Jia et al., UVB induces apoptosis via downregulation of CALML3-dependent JNK1/2 and ERK1/2 pathways in cataract. Int J Mol Med 41, 3041–3050 (2018).

19. Q. Zhu et al., CaMK II Inhibition Attenuates ROS Dependent Necroptosis in Acinar Cells and Protects against Acute Pancreatitis in Mice. Oxid Med Cell Longev 2021, 4187398 (2021).

20. F. Chen et al., Mitochondrial dysfunction in pancreatic acinar cells: mechanisms and therapeutic strategies in acute pancreatitis. Front Immunol 15, 1503087 (2024).

21. A. B. Tam, E. L. Mercado, A. Hoffmann, M. Niwa, ER stress activates NF-kappaB by integrating functions of basal IKK activity, IRE1 and PERK. PLoS One 7, e45078 (2012).

22. V. Delcroix et al., Lacrimal Gland Epithelial Cells Shape Immune Responses through the Modulation of Inflammasomes and Lipid Metabolism. Int J Mol Sci 24 (2023).

23. X. Chen, C. Shi, M. He, S. Xiong, X. Xia, Endoplasmic reticulum stress: molecular mechanism and therapeutic targets. Signal Transduct Target Ther 8, 352 (2023).

24. J. A. Kropski, T. S. Blackwell, Endoplasmic reticulum stress in the pathogenesis of fibrotic disease. J Clin Invest 128, 64–73 (2018).

25. L. Fu et al., The involvement of aquaporin 5 in the inflammatory response of primary Sjogren’s syndrome dry eye: potential therapeutic targets exploration. Front Med (Lausanne) 11, 1439888 (2024).

26. S. Hu et al., Lacrimal gland homeostasis is maintained by the AQP5 pathway by attenuating endoplasmic reticulum stress inflammation in the lacrimal gland of AQP5 knockout mice. Mol Vis 27, 679–690 (2021).

27. X. Mao et al., Loss of tricellular tight junction tricellulin leads to hyposalivation in Sjogren’s syndrome. Int J Oral Sci 17, 22 (2025).

28. K. Tsubota, S. Hirai, L. S. King, P. Agre, N. Ishida, Defective cellular trafficking of lacrimal gland aquaporin-5 in Sjogren’s syndrome. Lancet 357, 688–689 (2001).

29. E. Evans et al., Direct interaction between Rab3D and the polymeric immunoglobulin receptor and trafficking through regulated secretory vesicles in lacrimal gland acinar cells. Am J Physiol Cell Physiol 294, C662–674 (2008).

30. M. C. Edman et al., Increased Cathepsin S activity associated with decreased protease inhibitory capacity contributes to altered tear proteins in Sjogren’s Syndrome patients. Sci Rep 8, 11044 (2018).

31. E. Toth-Molnar, C. Ding, New insight into lacrimal gland function: Role of the duct epithelium in tear secretion. Ocul Surf 18, 595–603 (2020).

32. M. A. Shatos, L. Haugaard-Kedstrom, R. R. Hodges, D. A. Dartt, Isolation and characterization of progenitor cells in uninjured, adult rat lacrimal gland. Invest Ophthalmol Vis Sci 53, 2749–2759 (2012).

33. A. M. Chibly et al., Neurotrophin signaling is a central mechanism of salivary dysfunction after irradiation that disrupts myoepithelial cells. NPJ Regen Med 8, 17 (2023).

34. H. P. Makarenkova, D. A. Dartt, Myoepithelial Cells: Their Origin and Function in Lacrimal Gland Morphogenesis, Homeostasis, and Repair. Curr Mol Biol Rep 1, 115–123 (2015).

35. M. C. Adriance, J. L. Inman, O. W. Petersen, M. J. Bissell, Myoepithelial cells: good fences make good neighbors. Breast Cancer Res 7, 190–197 (2005).

36. V. Tepavcevic, R. R. Hodges, D. Zoukhri, D. A. Dartt, Signal transduction pathways used by EGF to stimulate protein secretion in rat lacrimal gland. Invest Ophthalmol Vis Sci 44, 1075–1081 (2003).

37. O. Mauduit et al., A closer look into the cellular and molecular biology of myoepithelial cells across various exocrine glands. Ocul Surf 31, 63–80 (2024).

38. L. Tedesco et al., An original amino acid formula favours in vitro corneal epithelial wound healing by promoting Fn1, ITGB1, and PGC-1alpha expression. Exp Eye Res 219, 109060 (2022).

39. V. L. Kolachala et al., Epithelial-derived fibronectin expression, signaling, and function in intestinal inflammation. J Biol Chem 282, 32965–32973 (2007).

40. M. Ponticos et al., Col1a2 enhancer regulates collagen activity during development and in adult tissue repair. Matrix Biol 22, 619–628 (2004).

41. S. S. Mathew-Steiner, S. Roy, C. K. Sen, Collagen in Wound Healing. Bioengineering (Basel) 8 (2021).

42. M. Sun et al., Collagen XIV Is an Intrinsic Regulator of Corneal Stromal Structure and Function. Am J Pathol 191, 2184–2194 (2021).

43. B. B. Young, G. Zhang, M. Koch, D. E. Birk, The roles of types XII and XIV collagen in fibrillogenesis and matrix assembly in the developing cornea. J Cell Biochem 87, 208–220 (2002).

44. L. Farr et al., CD74 Signaling Links Inflammation to Intestinal Epithelial Cell Regeneration and Promotes Mucosal Healing. Cell Mol Gastroenterol Hepatol 10, 101–112 (2020).

45. Y. Zhou et al., CD74 Deficiency Mitigates Systemic Lupus Erythematosus-like Autoimmunity and Pathological Findings in Mice. J Immunol 198, 2568–2577 (2017).

46. H. Su, N. Na, X. Zhang, Y. Zhao, The biological function and significance of CD74 in immune diseases. Inflamm Res 66, 209–216 (2017).

47. K. H. Kim et al., PRMT5 links lipid metabolism to contractile function of skeletal muscles. EMBO Rep 24, e57306 (2023).

48. H. Yoon, J. L. Shaw, M. C. Haigis, A. Greka, Lipid metabolism in sickness and in health: Emerging regulators of lipotoxicity. Mol Cell 81, 3708–3730 (2021).

49. L. Garcia-Posadas et al., Lacrimal Gland Myoepithelial Cells Are Altered in a Mouse Model of Dry Eye Disease. Am J Pathol 190, 2067–2079 (2020).

50. Z. She et al., The Role of B1 Cells in Systemic Lupus Erythematosus. Front Immunol 13, 814857 (2022).

51. J. Zikherman, K. Doan, R. Parameswaran, W. Raschke, A. Weiss, Quantitative differences in CD45 expression unmask functions for CD45 in B-cell development, tolerance, and survival. Proc Natl Acad Sci U S A 109, E3–12 (2012).

52. E. Montecino-Rodriguez, H. Leathers, K. Dorshkind, Identification of a B-1 B cell-specified progenitor. Nat Immunol 7, 293–301 (2006).

53. L. Y. Yim, C. S. Lau, V. S. Chan, Heightened TLR7/9-Induced IL-10 and CXCL13 Production with Dysregulated NF-ҝB Activation in CD11c(hi)CD11b(+) Dendritic Cells in NZB/W F1 Mice. Int J Mol Sci 20 (2019).

54. H. S. Carlsen, E. S. Baekkevold, H. C. Morton, G. Haraldsen, P. Brandtzaeg, Monocytelike and mature macrophages produce CXCL13 (B cell-attracting chemokine 1) in inflammatory lesions with lymphoid neogenesis. Blood 104, 3021–3027 (2004).

55. Z. Pan, T. Zhu, Y. Liu, N. Zhang, Role of the CXCL13/CXCR5 Axis in Autoimmune Diseases. Front Immunol 13, 850998 (2022).

56. T. Arkatkar et al., B cell-derived IL-6 initiates spontaneous germinal center formation during systemic autoimmunity. J Exp Med 214, 3207–3217 (2017).

57. O. Berczeli et al., Novel Insight Into the Role of CFTR in Lacrimal Gland Duct Function in Mice. Invest Ophthalmol Vis Sci 59, 54–62 (2018).

58. J. E. Wosen, D. Mukhopadhyay, C. Macaubas, E. D. Mellins, Epithelial MHC Class II Expression and Its Role in Antigen Presentation in the Gastrointestinal and Respiratory Tracts. Front Immunol 9, 2144 (2018).

59. M. J. Barrera et al., Endoplasmic reticulum stress in autoimmune diseases: Can altered protein quality control and/or unfolded protein response contribute to autoimmunity? A critical review on Sjogren’s syndrome. Autoimmun Rev 17, 796–808 (2018).

60. G. V. Cavalcanti et al., Correction: Endoplasmic reticulum stress in the salivary glands of patients with primary and associated Sjogren’s disease, and non-Sjogren’s sicca syndrome: a comparative analysis and the influence of chloroquine. Adv Rheumatol 65, 3 (2025).

61. E. Apostolou, P. Moustardas, T. Iwawaki, A. G. Tzioufas, G. Spyrou, Ablation of the Chaperone Protein ERdj5 Results in a Sjogren’s Syndrome-Like Phenotype in Mice, Consistent With an Upregulated Unfolded Protein Response in Human Patients. Front Immunol 10, 506 (2019).

62. W. Li et al., NF-kappaB and its crosstalk with endoplasmic reticulum stress in atherosclerosis. Front Cardiovasc Med 9, 988266 (2022).

63. V. R. Wiersma, M. Michalak, T. M. Abdullah, E. Bremer, P. Eggleton, Mechanisms of Translocation of ER Chaperones to the Cell Surface and Immunomodulatory Roles in Cancer and Autoimmunity. Front Oncol 5, 7 (2015).

64. E. V. Staikou et al., Calreticulin binds preferentially with B cell linear epitopes of Ro60 kD autoantigen, enhancing recognition by anti-Ro60 kD autoantibodies. Clin Exp Immunol 134, 143–150 (2003).

65. G. Kinoshita et al., Molecular chaperones are targets of autoimmunity in Ro(SS-A) immune mice. Clin Exp Immunol 115, 268–274 (1999).

66. J. G. Routsias, A. G. Tzioufas, M. Sakarellos-Daitsiotis, C. Sakarellos, H. M. Moutsopoulos, Calreticulin synthetic peptide analogues: anti-peptide antibodies in autoimmune rheumatic diseases. Clin Exp Immunol 91, 437–441 (1993).

67. T. Takeuchi, H. Ohno, IgA in human health and diseases: Potential regulator of commensal microbiota. Front Immunol 13, 1024330 (2022).

68. F. E. Johansen, C. S. Kaetzel, Regulation of the polymeric immunoglobulin receptor and IgA transport: new advances in environmental factors that stimulate pIgR expression and its role in mucosal immunity. Mucosal Immunol 4, 598–602 (2011).

69. M. Terzulli, L. Contreras-Ruiz, A. Kugadas, S. Masli, M. Gadjeva, TSP-1 Deficiency Alters Ocular Microbiota: Implications for Sjogren’s Syndrome Pathogenesis. J Ocul Pharmacol Ther 31, 413–418 (2015).

70. M. Wang, J. Huang, M. Lu, S. Zhang, C. Ding, ENaC in the Rabbit Lacrimal Gland and its Changes During Sjogren Syndrome and Pregnancy. Eye Contact Lens 41, 297–303 (2015).

71. P. Nandoskar et al., Changes of chloride channels in the lacrimal glands of a rabbit model of Sjogren syndrome. Cornea 31, 273–279 (2012).

72. J. Staal, L. P. Blanco, A. Perl, Editorial: Mitochondrial dysfunction in inflammation and autoimmunity. Front Immunol 14, 1304315 (2023).

73. M. A. Beckley, S. Shrestha, K. K. Singh, M. A. Portman, The role of mitochondria in the pathogenesis of Kawasaki disease. Front Immunol 13, 1017401 (2022).

74. A. Lepelley, T. Wai, Y. J. Crow, Mitochondrial Nucleic Acid as a Driver of Pathogenic Type I Interferon Induction in Mendelian Disease. Front Immunol 12, 729763 (2021).

75. J. W. Putney, G. S. Bird, Calcium signaling in lacrimal glands. Cell Calcium 55, 290–296 (2014).

76. S. Guo et al., Prostaglandin F2alpha exacerbated dry eye by promoting lacrimal gland fibrosis progression through the activation of the RhoA/ROCKs signaling pathway. Ocul Surf 38, 155–169 (2025).

77. M. Izumi et al., MR features of the lacrimal gland in Sjogren’s syndrome. AJR Am J Roentgenol 170, 1661–1666 (1998).

78. J. S. Pepose, R. F. Akata, S. C. Pflugfelder, W. Voigt, Mononuclear cell phenotypes and immunoglobulin gene rearrangements in lacrimal gland biopsies from patients with Sjogren’s syndrome. Ophthalmology 97, 1599–1605 (1990).

79. O. Mauduit et al., Spatial transcriptomics of the lacrimal gland features macrophage activity and epithelium metabolism as key alterations during chronic inflammation. Front Immunol 13, 1011125 (2022).

80. B. Parkin et al., Lymphocytic infiltration and enlargement of the lacrimal glands: a new subtype of primary Sjogren’s syndrome? Ophthalmology 112, 2040–2047 (2005).

81. M. Y. Wang, Y. Qiao, S. J. Wei, Z. L. Su, H. Y. Lu, MHC class II of different nonprofessional antigen-presenting cells mediate multiple effects of crosstalk with CD4(+)T cells in lung diseases. Front Med (Lausanne) 12, 1388814 (2025).

82. E. Grebenciucova, S. VanHaerents, Interleukin 6: at the interface of human health and disease. Front Immunol 14, 1255533 (2023).

83. R. C. Team (2021) R: A Language and Environment for Statistical Computing. (R Foundation for Statistical Computing, Vienna, Austria).

84. M. I. Love, W. Huber, S. Anders, Moderated estimation of fold change and dispersion for RNA-seq data with DESeq2. Genome Biol 15, 550 (2014).

85. J. Storey, The Positive False Discovery Rate: A Baysian Interpretation and the Q-Value. The Annals of Statistics 31, 2013–2035 (2003).

86. S. Xu et al., Using clusterProfiler to characterize multiomics data. Nat Protoc 19, 3292–3320 (2024).

87. X. Wang, J. Park, K. Susztak, N. R. Zhang, M. Li, Bulk tissue cell type deconvolution with multi-subject single-cell expression reference. Nat Commun 10, 380 (2019).

88. S. Jin et al., Inference and analysis of cell-cell communication using CellChat. Nat Commun 12, 1088 (2021).

89. B. Diedenhofen, J. Musch, cocor: a comprehensive solution for the statistical comparison of correlations. PLoS One 10, e0121945 (2015).

90. H. Wickham, ggplot2: Elegant Graphics for Data Analysis (Springer-Verlag New York, 2016).

91. A. Kassambara (2025) ggpubr: ‘ggplot2’ Based Publication Ready Plots.

92. R. Kolde (2019) pheatmap: Pretty Heatmaps.

