## Supplement Table-1 and Figures 1-4 for "Spatial transcriptomic profiling identifies lacrimal gland epithelial cell-driven mechanisms underlying autoimmunity in Sjögren’s disease"

**Supplementary Table 1.** Antibodies Used.

| Target | Host Species/Conjugate | Clone | Vendor (Catalogue #) |
|---|---|---|---|
| **Primary Antibodies** | | | |
| **B220** | Rat (IgG2a), PE-Cy5 | RA3-6B2 | BD Biosciences (553091) |
| **B220** | Rat (IgG2a) | RA3-6B2 | Biolegend (103202) |
| **BCL6** | Rabbit (IgG) | Polyclonal | Fabgennix (BCL6-601AP) |
| **Calreticulin** | Rabbit (IgG) | SU37-03 | Invitrogen (MA5-32131) |
| **CFTR** | Mouse (IgG1) | A3 | Santa Cruz (SC-376683) |
| **CXCR5** | Rabbit (IgG) | JB11-40 | Invitrogen (MA5-41258) |
| **Cytokeratin** | Mouse (IgG) | AE1/AE3 | MP Biomed (08691451) |
| **a-ENAC** | Rabbit (IgG) | HL3027 | Invitrogen (MA5-56694) |
| **Rab3D** | Goat (IgG) | Polyclonal | Santa Cruz (SC-26392) |
| **a -SMA** | Mouse (IgG2a) | 1A4 | Invitrogen (18-0106) |
| **CD45** | Rat (IgG2b), AF 488 | 30-F11 | Biolegend (103121) |
| **Secondary Antibodies** | | | |
| **Rabbit IgG** | Goat (IgG), AF 568 | Polyclonal | Invitrogen (A-11011) |
| **Rabbit IgG** | Goat (IgG), AF 488 | Polyclonal | Invitrogen (A-11008) |
| **Rat IgG** | Goat (IgG), AF 568 | Polyclonal | Invitrogen (A-11077) |
| **Mouse IgG** | Goat (IgG), AF 568 | Polyclonal | Invitrogen (A-11004) |
| **Goat IgG** | Donkey (IgG), AF568 | Polyclonal | Invitrogen (A-11057) |

### Supplementary Figure1

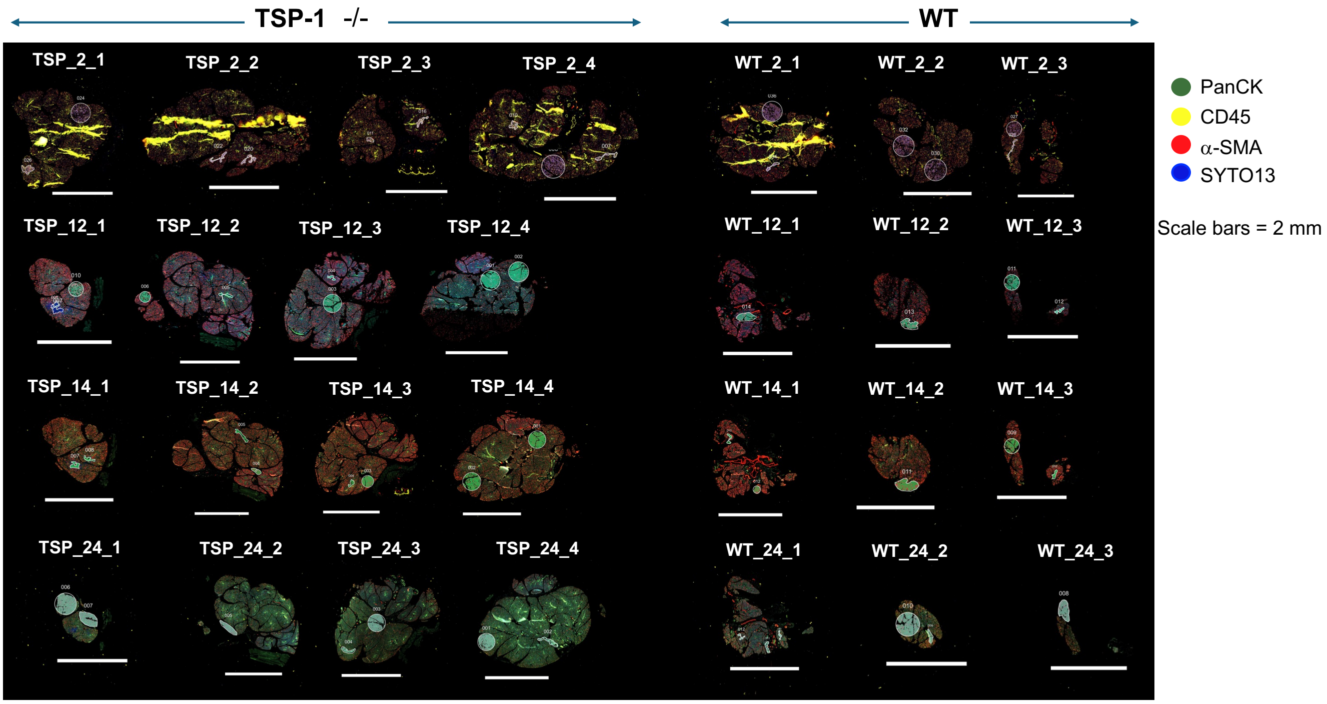


**S1: Digital spatial profiling with whole transcriptome atlas (WTA DSP) of mouse lacrimal glands.** Immunofluorescence images of FFPE sections from WT and TSP-1$^{-/-}$ LGs stained with morphology markers with each section analyzed for the transcriptome of marked ROIs. Color legend indicates target for each fluorophore-conjugated antibody used as morphology marker to identify cell types – epithelial cells (PanCK), Immune cells (CD45), Myoepithelial cells (a-SMA) and nuclear stain (SYTO 13). .

**Supplementary Figure 2**

Acinar epithelial cells

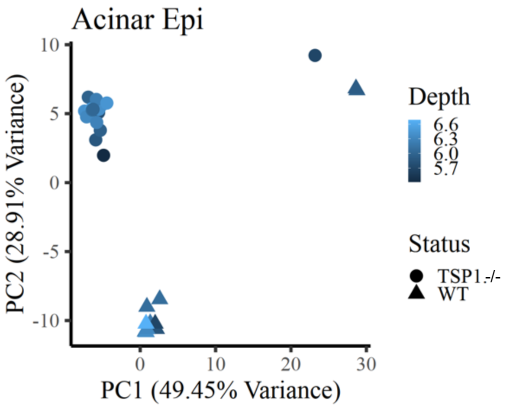


Duct epithelial cells

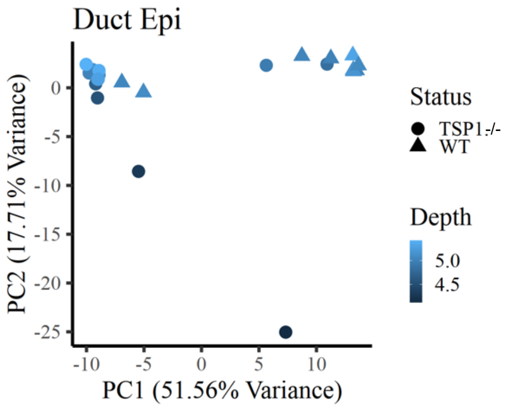


Myoepithelial cells

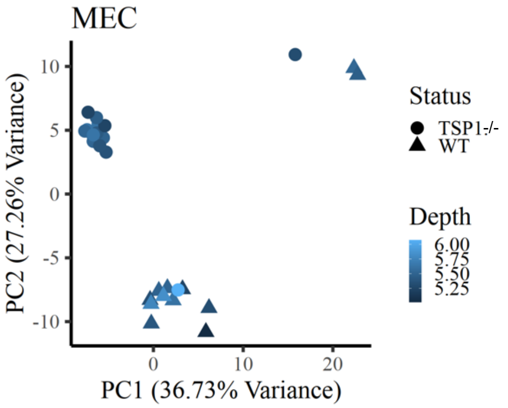


**S2: Epithelial cell clustering in analyzed ROIs**
Principal component analysis plots showing epithelial cell clusters detected in ROIs marked in WT and TSP-1 deficient LG sections.

### Supplementary Figure 3

**A.**

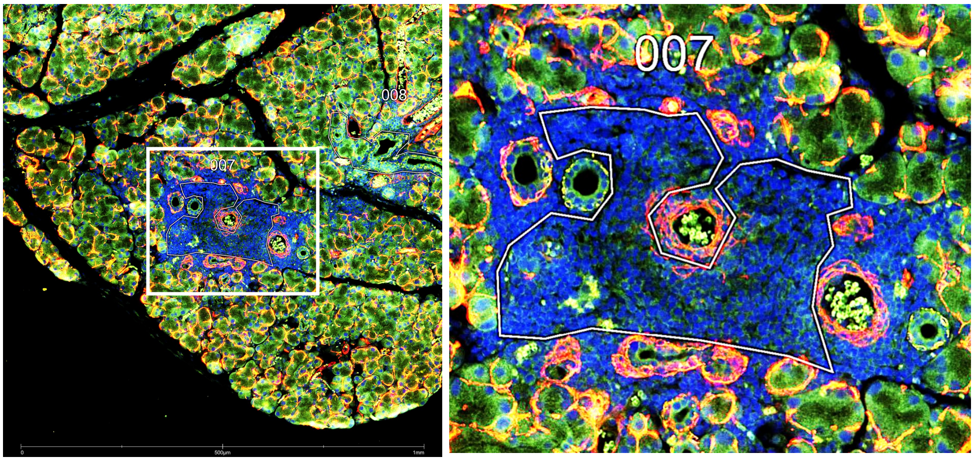


**B.**

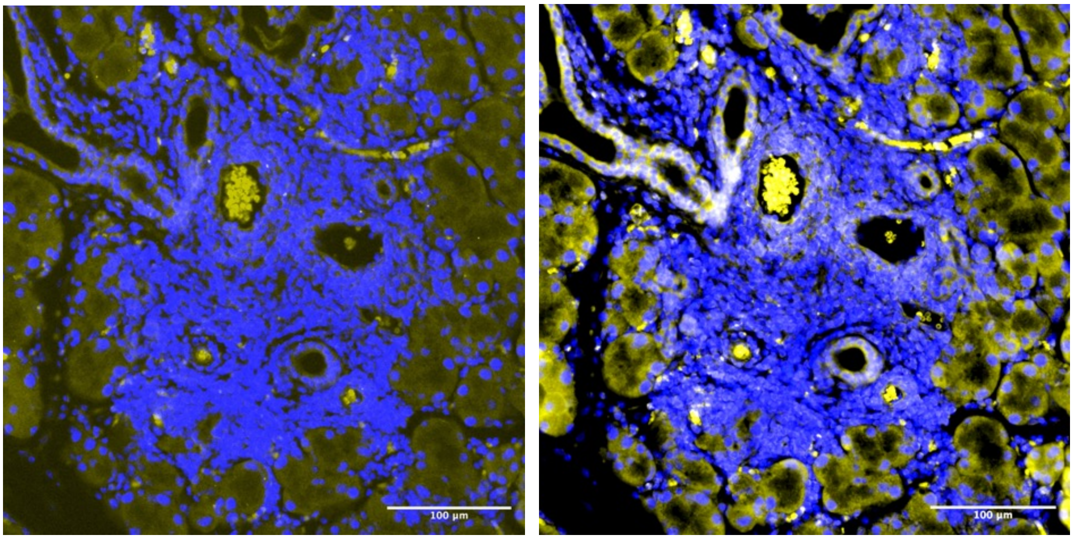


**S3: Weak CD45 staining of immune infiltrates in TSP-1 deficient lacrimal gland.**
A. Lacrimal gland tissue section from TSP-$1^{-/-}$ mouse showing infiltrate containing ROI stained weakly (yellow) with morphology marker anti-CD45 (clone EM-05 and fluorescent secondary antibody) used in DSP analysis. Immune infiltrates are detectable with blue nuclear staining within marked ROI; B. Confirmation of weak CD45 staining (yellow) pattern within immune infiltrates in TSP-$1^{-/-}$ LG section stained with fluorescence-conjugated anti-CD45 (clone 30-F11) (right), control antibody (left) and DAPI for nuclear staining.

Supplementary Figure 4

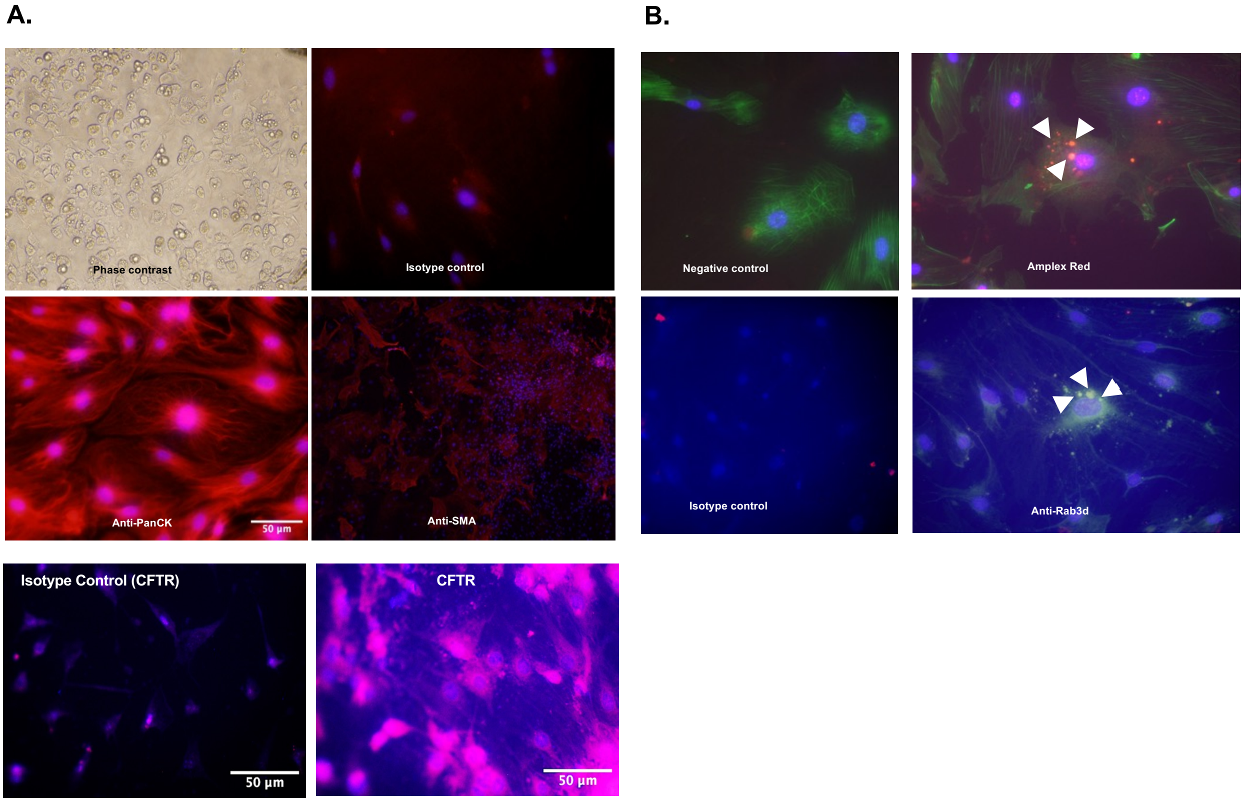


**S4: Primary cultures of lacrimal gland epithelial cells**

A. Adherent primary cultures stained predominantly with epithelial cell marker (PanCK) and duct epithelial marker (CFTR) with negligible (< 5%) staining for myoepithelial marker (SMA); B. Primary cultures immunostained with Phalloidin (green) and Amplex Red reagent (to stain secretory vesicles containing peroxidase) or green fluorochrome conjugated anti-Rab3d (secretory vesicle marker); arrowheads indicate positively stained secretory vesicles of acinar epithelial cells (< 5% of total cells). Nulear stain DAPI (Blue). Images at 200x magnification.
